# TORC1 regulates vacuole membrane composition through ubiquitin- and ESCRT-dependent microautophagy

**DOI:** 10.1101/854760

**Authors:** Xi Yang, Weichao Zhang, Xin Wen, Patrick J. Bulinski, Dominic A. Chomchai, Felichi Mae Arines, Yun-Yu Liu, Simon Sprenger, David Teis, Daniel J. Klionsky, Ming Li

## Abstract

Cellular adaptation in response to nutrient limitation requires the induction of autophagy and lysosome biogenesis for the efficient recycling of macromolecules. Here, we discovered that starvation and TORC1 inactivation not only lead to the upregulation of autophagy and vacuole proteins involved in recycling, but also result in the downregulation of many vacuole membrane proteins to supply amino acids as part of a vacuole remodeling process. Downregulation of vacuole membrane proteins is initiated by ubiquitination, which is accomplished by the coordination of multiple E3 ubiquitin ligases, including Rsp5, the Dsc complex, and a newly characterized E3 ligase, Pib1. The Dsc complex is negatively regulated by TORC1 through the Rim15-Ume6 signaling cascade. After ubiquitination, vacuole membrane proteins are sorted into the lumen for degradation by ESCRT-dependent microautophagy. Thus, our study uncovered a complex relationship between TORC1 inactivation and vacuole biogenesis.

## Introduction

The lysosome is a central catabolic station that breaks down and recycles cellular materials to maintain nutrient homeostasis (Lim and Zoncu, 2016; Settembre et al., 2013). Emerging evidence suggests that the lysosome is a dynamic organelle, which constantly adjusts its membrane composition according to environmental cues. In yeast, changes in substrate concentration can lead to the degradation of their corresponding vacuole (yeast lysosome) membrane transporters. For example, depleting lysine from the cytoplasm triggers a selective degradation of the vacuolar lysine importer Ypq1 to stop lysine from being transported from the cytoplasm into the vacuole lumen (Li et al., 2015b; Sekito et al., 2014). Similarly, a low cytoplasmic Zn^2+^ level leads to the downregulation of a vacuolar Zn^2+^ importer, Cot1 (Li et al., 2015a).

Our understanding of the underlying mechanisms for vacuole membrane regulation is still incomplete. However, it has been shown that protein ubiquitination serves as a sorting signal to initialize the degradation process. In yeast, two independent vacuole E3 ligases are involved in the substrate-triggered degradation of transporters. Specifically, a cytosolic NEDD4 family E3 ligase, Rsp5, and its vacuole membrane adapter, Ssh4, are responsible for the ubiquitination of Ypq1 (Li et al., 2015b), whereas Dsc, a multi-subunit transmembrane ubiquitin ligase complex, is required for the ubiquitination of Cot1 (Li et al., 2015a; Yang et al., 2018). After ubiquitination, vacuole membrane proteins are delivered into the lumen by the endosomal sorting complexes required for transport (ESCRT) pathway (Li et al., 2015a; Li et al., 2015b; Zhu et al., 2017). With this selective degradation mechanism, the vacuole membrane composition is regulated accurately in response to specific transporting substrates. Besides this fine level adjustment of individual transporters, how do other environmental stresses such as starvation regulate vacuole membrane composition?

Beyond catabolism, the lysosome also plays a major role in cellular stress response. The evolutionarily conserved mechanistic target of rapamycin kinase complex 1 (MTORC1), a lysosome membrane-associated kinase complex, serves as the signaling hub to sense different stresses and regulate cellular metabolism accordingly (Laplante and Sabatini, 2009; Perera and Zoncu, 2016). In mammalian cells, under nutrient-rich conditions, MTORC1 is recruited to the lysosome membrane and activated to phosphorylate the ribosomal RPS6 kinases (RPS6KB1/S6K1, RPS6KB2/S6K2) and ElF4EBP1 (eukaryotic translation initiation factor 4E binding protein 1) to promote protein synthesis, leading to cell growth and proliferation (Holz et al., 2005). In addition, active MTORC1 inhibits the transcription of various stress response genes involved in autophagy, lysosome biogenesis, and endoplasmic reticulum (ER) stress by phosphorylating TFEB (transcription factor EB) and TFE3 (Rehli et al., 1999; Sardiello et al., 2009; Settembre et al., 2011; Settembre et al., 2012). Similarly, in yeast, active TORC1 directly phosphorylates Sch9, an analog of the mammalian RPS6 kinases, to stimulate protein translation when nutrients are available (Jin et al., 2014; Urban et al., 2007). Meanwhile, TORC1 also inhibits autophagy through Atg13 phosphorylation, thereby inhibiting its ability to activate the Atg1 kinase (Fujioka et al., 2014; Kraft et al., 2012).

Conversely, when TORC1/MTORC1 is inactive under stress conditions, catabolic processes such as proteasome degradation, autophagy, and endocytosis of plasma membrane proteins are elevated. In addition, lysosome biogenesis is upregulated to boost its degradative and recycling function. In mammalian cells, when MTORC1 is inactive, dephosphorylated TFEB translocates into the nucleus to promote the transcription of numerous target genes, including those encoding lysosomal hydrolases, pumps, permeases, and lysosome positioning regulators (Martina et al., 2014; Puertollano et al., 2018; Sardiello et al., 2009; Settembre et al., 2011; Settembre et al., 2012; Willett et al., 2017). In yeast, many vacuolar proteases such as Prb1, Ape1, Cps1, and Pep4 are upregulated to enhance the digestion function during starvation (Muller et al., 2015; Parzych and Klionsky, 2018). All these studies point to a direct correlation between MTORC1/TORC1 inactivation and the enhancement of lysosomal/vacuolar biogenesis. However, two recent studies suggested the complexity of their relationship in yeast. Sakai and colleagues observed that glucose depletion leads to the invagination and degradation of two vacuole membrane proteins, Vph1 (a V_0_ subunit of the vacuolar ATPase) and Pho8 (vacuolar phosphatase), in an ESCRT-dependent manner (Oku et al., 2017). Similarly, De Virgilio and colleagues observed that rapamycin treatment leads to the ESCRT-dependent degradation of Pho8 (Hatakeyama et al., 2019). These observations indicated that remodeling the vacuole proteome after TORC1 inactivation might be more complex than previously anticipated, and raised several interesting questions: First, is the downregulation of vacuole membrane proteins a general response to TORC1 inactivation? Second, what machinery is involved? And third, how does TORC1 regulate this machinery?

In this study, we report that TORC1 plays a critical role in regulating vacuole membrane composition. In contrast to the simplified model that TORC1 inactivation leads to a global upregulation of vacuole biogenesis, we discovered that TORC1 inactivation also triggers the concomitant downregulation of numerous vacuole membrane proteins. Further analysis revealed that multiple E3 ligase systems, including Rsp5, the Dsc complex, and a third E3 ligase, Pib1, function together on the vacuole membrane to ubiquitinate proteins. Moreover, our results showed that TORC1 can regulate the activity of the vacuole ubiquitin machinery. Specifically, TORC1 regulates the production and assembly of the vacuolar Dsc complex. After ubiquitination, cargoes are sorted into the lumen for degradation in an ESCRT-dependent manner. This study thus extends our understanding of the complexity of how TORC1 regulates vacuole composition according to environmental cues.

## Results

### TORC1 inactivation triggers downregulation of many vacuole membrane proteins

The TORC1 complex is responsible for regulating numerous stress responses according to environmental cues. However, how TORC1 regulates the vacuole protein composition is only partially understood. To answer this question, we set out to understand how vacuole membrane composition responds to environmental stresses such as starvation, and its relationship with TORC1 inactivation. As an initial test, we monitored the levels of five vacuole membrane proteins, including Vba4 (a putative amino acid permease), Fet5 (a subunit of the putative iron transporter complex, which dimerizes with Fth1), Fth1, Vph1, and Zrt3* (a variant of the zinc exporter Zrt3, please see Li et al., 2015a for details), after yeast cells enter the stationary phase. All five vacuole membrane proteins were chromosomally tagged with the green fluorescent protein (GFP), and they properly localized to the vacuole membrane (Fig. S1A) (Li et al., 2015a). We then used a GFP antibody to measure the protein level changes. As shown in Fig. 1A-B, we took four time points (16, 20, 24, and 36 h) in the stationary phase and compared them to mid-log cells (1 h, with OD_600_ ∼0.7). All five proteins were significantly downregulated after cells entered the stationary phase. During degradation, free GFP accumulated due to its resistance to vacuolar proteases (Fig. 1A-B and S1A). Longer incubation in the stationary phase led to a higher level of protein degradation and more accumulation of free GFP (Fig. 1A, 16-36 h). In contrast, very little degradation was observed in mid-log cells (Fig. 1A, 1 h). In addition to this natural starvation, acute nitrogen starvation also caused a similar downregulation of these proteins (data not shown). Together, our results suggested that long-term nutrient starvation can trigger the downregulation of many vacuole membrane proteins.

**Figure 1.**
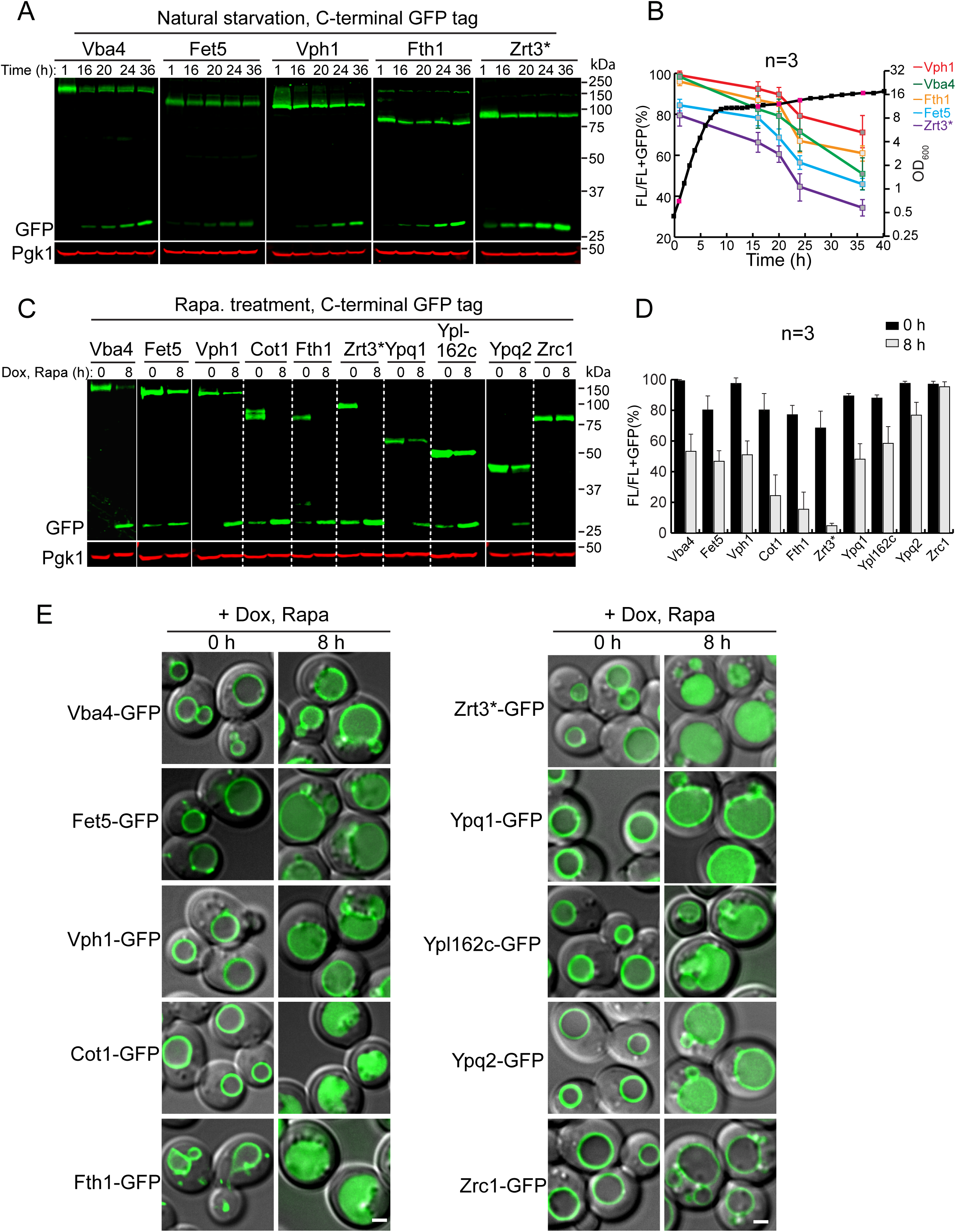
TORC1 inactivation triggers the downregulation of many vacuole membrane proteins. **(A)** Western blots showing the downregulation of five vacuole membrane proteins in stationary phase cells. Samples were collected at the indicated time points and 1 OD_600_ units of cells were loaded in each lane. **(B)** Quantification of the protein levels in (A). The black curve represents the yeast growth curve in YPD medium at 28°C, and the red squares on the growth curve represent the time points chosen for western blot analysis. FL: full-length protein fused with GFP. The relative protein levels were calculated as FL/(FL + free GFP). **(C)** Western blots showing the downregulation of TET-off-controlled vacuole membrane proteins after rapamycin treatment. The same volume of cells was loaded, with 0.5 OD_600_ units of cells loaded at 0 h. **(D)** Quantification of the protein levels in (C). **(E)** Merged images (DIC+GFP) to show subcellular localization of vacuole membrane proteins before (0 h) and after (8 h) rapamycin treatment. Scale bar: 2 μm.

The downregulation of vacuole membrane proteins in response to nutrient starvation is a surprising finding, especially because it has been widely assumed that starvation promotes vacuole biogenesis and autophagy to boost the recycling of intracellular materials (Noda, 2017). Consistently, previous studies have shown that vacuolar hydrolases, components of the autophagic machinery, and the vacuole membrane amino acid exporter Atg22, are indeed upregulated under starvation conditions (Muller et al., 2015; Yang et al., 2006).

To verify those reported observations, we collected antibodies against endogenous vacuolar/autophagic proteins, including Vph1, Pep4, Cps1, and Atg8, and checked their response to natural starvation. Due to the lack of an Atg22 antibody, we chromosomally tagged Atg22 with GFP at its C terminus. As shown in Fig. S1B-C, vacuolar proteases (Pep4 and Cps1) and autophagic machinery (Atg8 and Atg22) were indeed induced by starvation. In contrast, the protein level of untagged Vph1 decreased, which was consistent with the observation made in Fig. 1A where Vph1-GFP was partially degraded. Together, our data indicated that vacuoles undergo extensive remodeling in response to nutrient limitation. Proteins involved in the digestion and recycling functions are upregulated, while many other membrane proteins are downregulated. Interestingly, an accumulation of free GFP was also observed for Atg22-GFP (Fig. S1D), indicating that even a protein directly involved in recycling can still be subjected to degradation after long-term starvation.

To directly test if the downregulation is due to TORC1 inactivation, we treated the mid-log phase cells with rapamycin. In this experiment, we expanded the substrate list to ten GFP-tagged vacuole membrane proteins by adding Cot1, Ypq1, Ypl162c (a putative transporter with unknown function), Ypq2 (a homolog of Ypq1), and Zrc1 (another zinc importer, and a homolog of Cot1) (Fig. 1C-E). To focus on the pre-existing pool of vacuole membrane proteins, we expressed them under the control of a TET-OFF system (Gari et al., 1997) and pretreated yeast cells with doxycycline to stop their gene transcription before rapamycin treatment. In this assay, nine out of ten tested vacuole membrane proteins were downregulated to different levels, and the accumulation of a lumenal GFP signal was observed (Fig. 1C-E). The only exception was seen with Zrc1, which was unchanged after rapamycin treatment. This result suggested that maintaining the Zn^2+^ import capability is important for the recycling function of the vacuole, which contains many zinc-dependent metalloenzymes (Hecht et al., 2014; Hecht et al., 2013; Simm et al., 2007). Nevertheless, our results strongly suggested that TORC1 inactivation triggers the downregulation of many vacuole membrane proteins, which is in contrast to the long-held view that vacuole biogenesis will be globally upregulated after TORC1 inactivation.

### The degradation of vacuole membrane proteins depends on lumenal proteases

The accumulation of a lumenal GFP signal corresponding to the various chimeric reporters we tested (Fig. 1E) after rapamycin treatment indicated that the degradation happens inside the vacuole. To confirm this, we chose four vacuole membrane proteins (Ypl162c-GFP, Ypq1-GFP, Vph1-GFP, and Cot1-GFP) and performed the degradation assay in a *pep4*Δ strain. Pep4 is the master protease that processes and activates other vacuolar zymogens (Ammerer et al., 1986; Woolford et al., 1986). Deleting the *PEP4* gene results in general defects in the vacuolar protease activity. As shown in Fig. 2, for all four tested substrates, full-length proteins were stabilized in the *pep4*Δ strain, and the accumulation of free GFP was entirely abolished. In addition, we observed a band shift for Ypl162c-GFP in the *pep4*Δ strain (Fig. 2A), suggesting that after reaching the vacuole membrane, Ypl162c undergoes a proteolytic cleavage by lumenal proteases for its maturation. In conclusion, these results indicated that the degradation depends on vacuolar protease activities.

**Figure 2.**
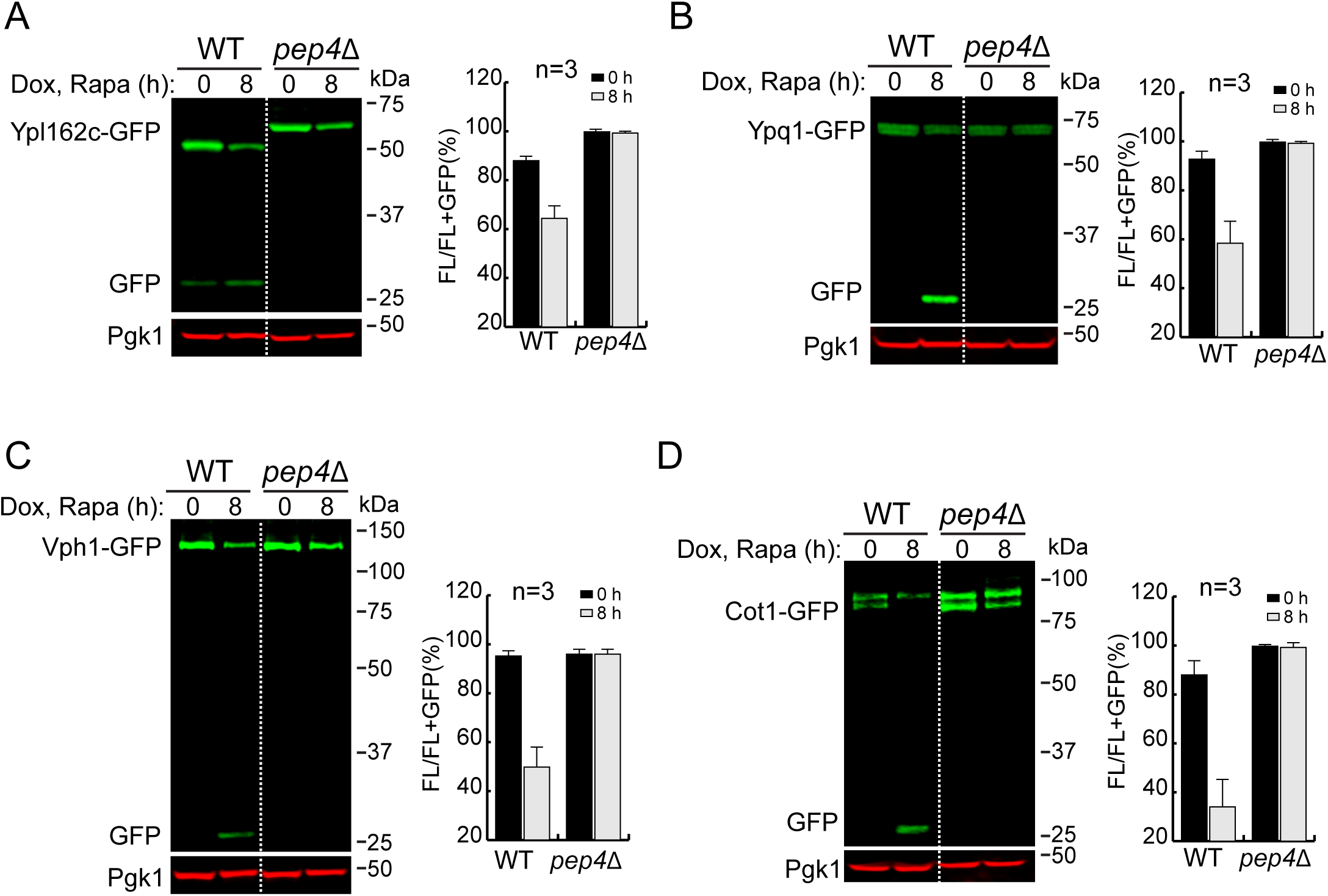
The degradation of vacuole membrane proteins depends on lumenal proteases. **(A-D)** Western blots (left) and corresponding quantifications (right) showing the degradation of (A) Ypl162c-GFP, (B) Ypq1-GFP, (C) Vph1-GFP or (D) Cot1-GFP in WT and *pep4*Δ strain cells. The same volume of cells was loaded, with 0.5 OD_600_ units of cells loaded at 0 h.

### The ESCRT machinery is required for invagination into the vacuole lumen

So far, three pathways have been suggested in yeast for delivering proteins to the vacuole lumen: macroautophagy, ESCRT-dependent sorting of ubiquitinated cargoes via the MVB pathway or microautophagy, and the ESCRT-independent intralumenal fragment (ILF) pathway. Among them, the ILF pathway proposes that the single vesicle (named as an intralumenal fragment) created by vacuole-vacuole homotypic fusion is responsible for selectively sorting membrane proteins into the lumen for degradation (McNally et al., 2017). Although the protein sorting mechanism has not been addressed, the ILF pathway has been reported to be blocked by rapamycin treatment and stimulated by cycloheximide-triggered hyperactivation of TORC1 (McNally et al., 2017), hence making it unlikely to be responsible for internalizing vacuole membrane proteins after TORC1 inactivation.

We then tested whether macroautophagy or ESCRT machinery were responsible for the degradation. Deleting *ATG1*, an essential gene for macroautophagy, had little effect on the degradation of all tested membrane substrates (Cot1-GFP, Vph1-GFP, Ypl162c-GFP, and Ypq1-GFP, Fig. S2A-D). In contrast, all the substrates were stabilized when two independent ESCRT components, *VPS4 (*the AAA-ATPase that disassembles the ESCRT-III filaments) and *VPS27* (a component of the ESCRT-0 subcomplex), were deleted (Fig. S2A-D and S3). These results showed that the ESCRT machinery, but not macroautophagy, is required for the degradation of vacuole membrane proteins (Hatakeyama et al., 2019; Oku et al., 2017; Zhu et al., 2017).

One caveat of using ESCRT deletion strains was the formation of class E compartments, which refers to the aberrant endosomal structures that are adjacent to vacuoles after the ESCRT machinery is inactivated. These aberrant endosomes can no longer efficiently fuse with the vacuole membrane. As such, most proteins that travel through the MVB pathway are trapped outside the vacuole, including Vph1-GFP (MacDonald et al., 2012; Yang et al., 2018; Zhu et al., 2017). As shown in Fig. S2E and S3E, in an ESCRT deletion strain, the majority of Vph1-GFP was trapped at the class E compartment. In contrast, membrane proteins that travel through the independent AP-3 pathway, including Ypl162c-GFP, Cot1-GFP, Ypq1-GFP, and Zrc1-mCherry, are mostly localized to the vacuole membrane (Fig. S2E and S3E). This observation raises the possibility that the block of Vph1-GFP degradation is because it cannot reach the vacuole, and thus cannot be recognized by the vacuole degradation machinery.

To exclude this possibility, we performed the degradation assay in a temperature-sensitive *vps4^ts^* strain (Babst et al., 1997). Yeast cells were first grown at the permissive temperature (26°C) to ensure all vacuole membrane proteins, including substrates and the required degradation machinery, traffick normally to the vacuole. Then the degradation assay was performed at both 26°C (permissive) and 37°C (non-permissive) temperatures. At 26°C, the degradation of all tested substrates occurred normally (Fig. 3A-E). However, after cells were shifted to 37°C, their degradation was drastically reduced, and proteins were stabilized on the vacuole membrane.

**Figure 3.**
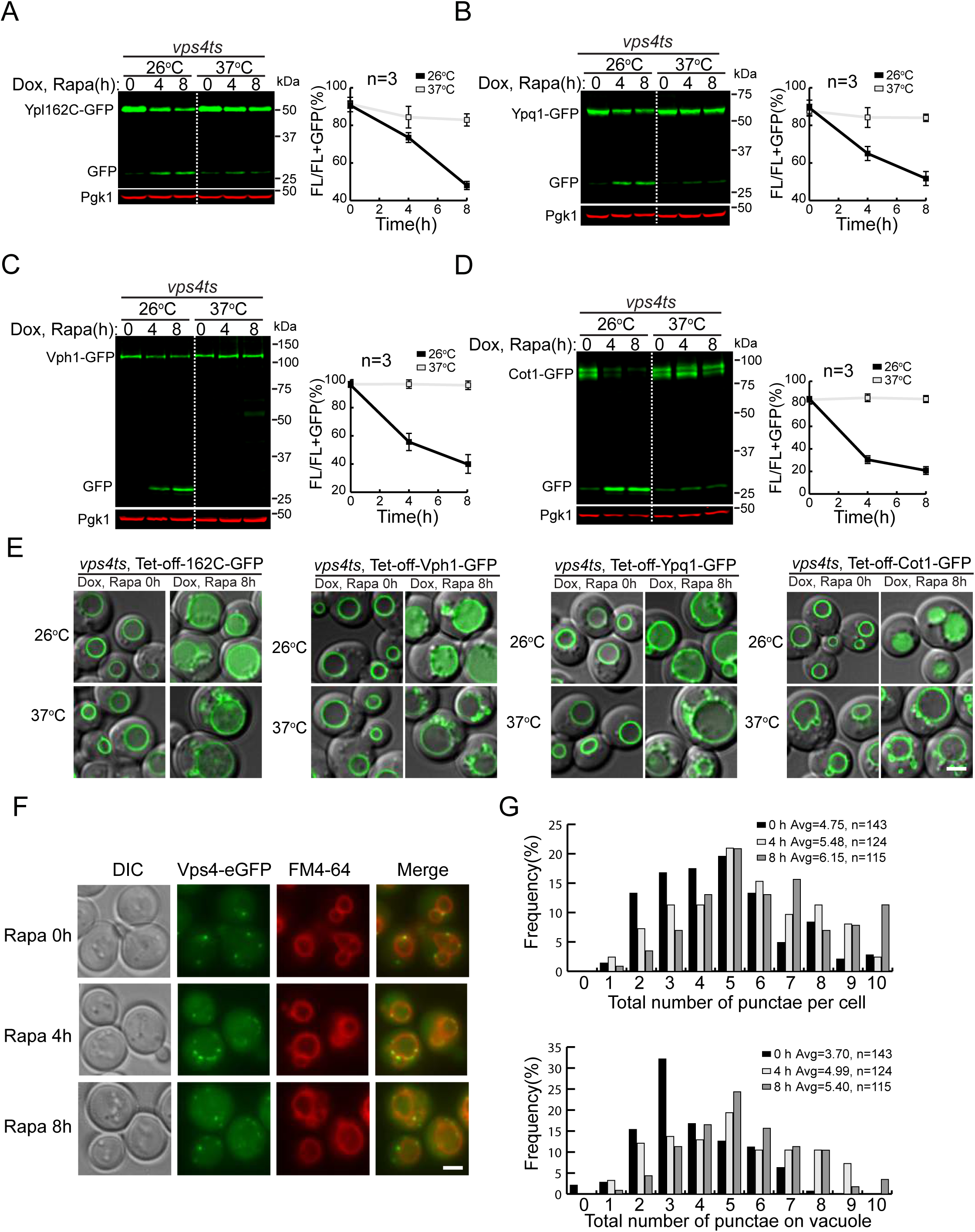
The ESCRT machinery is required for the degradation of vacuole membrane proteins. **(A-D)** Western blots (left) and corresponding quantifications (right) showing the degradation of (A) Ypl162c-GFP, (B) Ypq1-GFP, (C) Vph1-GFP or (D) Cot1-GFP in *vps4^ts^* cells at both 26°C and 37°C. The same volume of cells was loaded, with 0.5 OD_600_ units of cells loaded at 0 h. **(E)** Subcellular localization of Ypl162c-GFP, Vph1-GFP, Ypq1-GFP or Cot1-GFP in *vps4^ts^* cells at both 26°C and 37°C after rapamycin treatment. **(F)** Subcellular localization of Vps4-eGFP before (0 h) and after (4 h, 8 h) rapamycin treatment. **(G)** Quantification of the Vps4-eGFP puncta in (F). Scale bar: 2 μm.

If the ESCRT machinery is responsible for increased degradation of vacuole membrane proteins, one would expect to see more ESCRTs localize (or adjacent) to the vacuole membrane after TORC1 inactivation. To test this, we tagged Vps4 with a functional 3HA-eGFP tag (Vps4-3HA-eGFP)(Adell et al., 2017) and checked its localization upon TORC1 inactivation. As shown in Fig. 3F-G, after rapamycin treatment, more Vps4-GFP puncta were localized at or in direct vicinity of the vacuole membrane (4 h, 8 h vs. 0 h), supporting a model in which the ESCRT machinery was active at the membrane to sort cargoes.

Lastly, we used transmission electron microscopy to visualize the vacuole membrane invagination after 4 hours of rapamycin treatment. As shown in Fig. 4A-C, three types of invagination were observed in WT cells. The most prominent group was the <u>p</u>iecemeal <u>m</u>icroautophagy of the <u>n</u>ucleus (PMN, ∼26%, n=82, Fig. 4J). Besides PMN that happens at the nucleus-vacuole junctions, we also observed <u>cy</u>toplasmic <u>m</u>icroautophagy of <u>o</u>rganelles such as lipid droplets (CMO, ∼10%, n=82, Fig. 4K). The third group was the small tubular invagination of vacuole membrane (∼14.5%, n=82, Fig. 4A-B and inserts, 4G). In addition, we also observed small vesicles inside the vacuole (Fig 4A-F, arrows). After deleting *VPS27*, the small tubular microautophagy was nearly abolished (∼1.2%, Fig. 4D-F, 4G, n=74). Moreover, the number of small vesicles inside the vacuole was drastically reduced (Fig. 4H). In contrast, the other two forms of microautophagy was either unaffected (for CMO) or reduced but not abolished (for PMN). Combined with the Vps4 localization analysis, these data supported a model that the ESCRT localizes to the vacuole membrane to directly invaginate cargo proteins as small vesicles (Fig. 4A-C, 4I).

**Figure 4.**
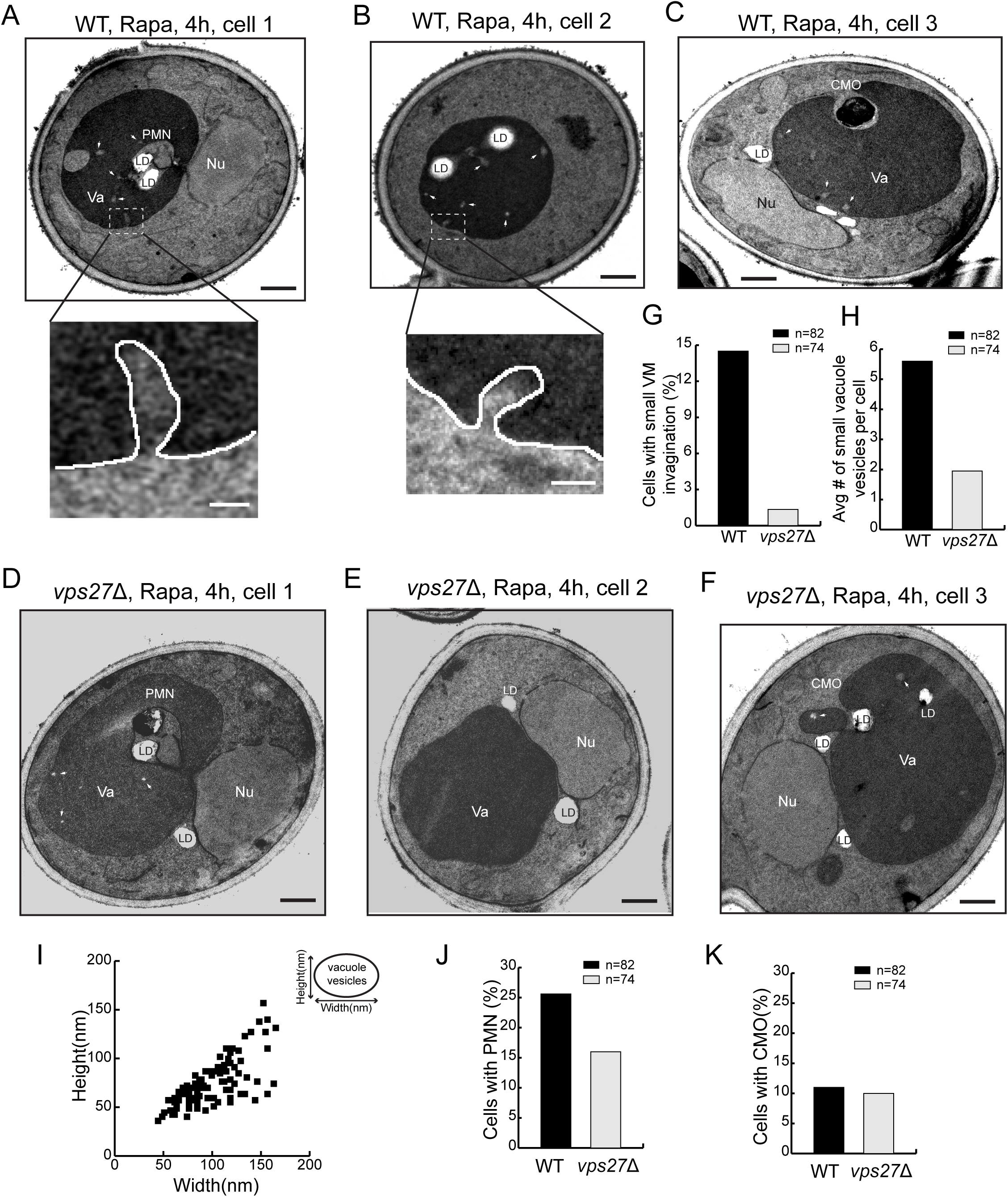
The ESCRT deletion abolished one form of microautophagy. **(A-C)** Representative TEM images showing three types of microautophagy, including piecemeal microautophagy of the nucleus (PMN), cytoplasmic microautophagy of organelles (CMO), and small vacuole membrane (VM) invagination in WT cells after rapamycin treatment. **(D-E)** Representative TEM images showing that after rapamycin treatment, PMN and CMO still happened in *vps27*Δ cells, while the small VM invagination was nearly abolished. **(G)** Frequency of observing small VM invagination in either WT or *vps27*Δ cells after rapamycin treatment. **(H)** Number of small vacuole vesicles per cell in either WT or *vps27*Δ cells after rapamycin treatment**. (I)** Size distribution of small vacuole vesicles. **(J)** Frequency of observing PMN in either WT or *vps27*Δ cells after rapamycin treatment. **(K)** Frequency of observing CMO in either WT or *vps27*Δ cells after rapamycin treatment. Va: Vacuole; Nu: Nucleus; LD: Lipid droplet. White arrows highlight small vacuole vesicles. Black scale bar: 0.5 μm. White scale bar: 50 nm.

Taken together, our results indicated that the ESCRT-mediated microautophagy, but not ILF or macroautophagy, is essential for the sorting of vacuole membrane proteins into the lumen.

### Multiple E3 ubiquitin ligases function at the vacuole

Ubiquitination is a prerequisite for cargo recognition by the ESCRT machinery, which implies that E3 ligases are important for the starvation-triggered vacuole membrane degradation. As of now, two independent E3 ligase complexes, the Ssh4-Rsp5 complex and the Dsc complex, have been identified to ubiquitinate vacuole membrane proteins (Li et al., 2015a; Li et al., 2015b; Yang et al., 2018). However, only three vacuole transporters (Ypq1, Cot1, and Zrt3*) have been shown to be their substrates. Our observation that rapamycin treatment triggers the downregulation of many vacuole membrane proteins expands their potential substrate repertoires.

To evaluate the importance of these E3 ligases in TORC1-mediated downregulation, we performed degradation assays in wild-type (WT), *ssh4*Δ, *tul1*Δ (the E3 ligase in the Dsc complex), and *ssh4*Δ *tul1*Δ double-mutant strains (Fig. 5 and S4). Interestingly, our tested substrates showed a diverse E3 ligase preference. Based on the E3 ligase dependence, we divided them into three groups: (A) Ssh4-dependent, (B) Tul1- and unknown E3 ligase-dependent, and (C) Ssh4-, Tul1-, and unknown E3 ligase-dependent.

**Figure 5.**
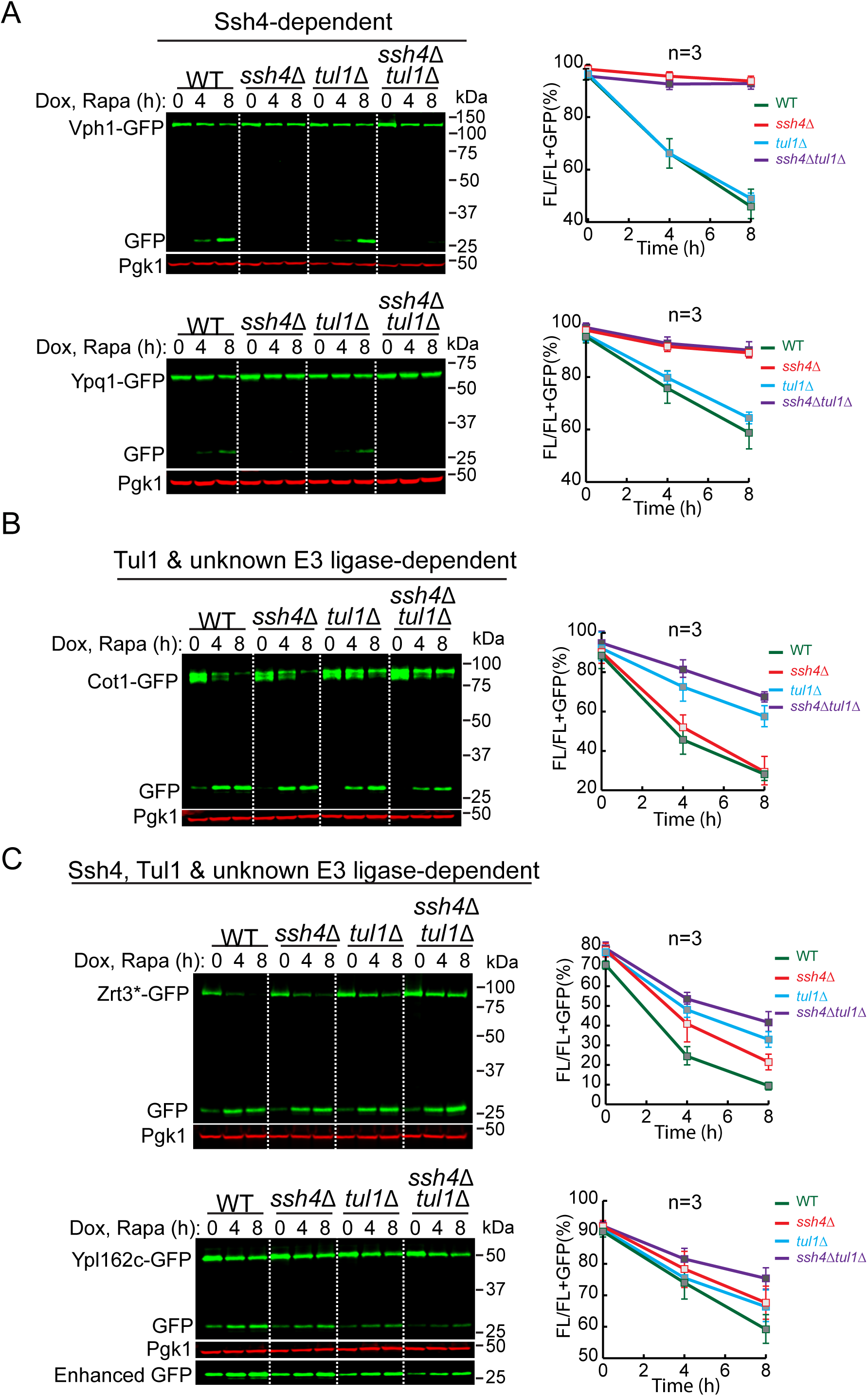
Multiple vacuole E3 ligases function downstream of the TORC1 kinase. **(A-C)** Western blots (left) and corresponding quantifications (right) showing the degradation of (A) Vph1-GFP and Ypq1-GFP, (B) Cot1-GFP, or (C) Zrt3*-GFP and Ypl162c-GFP in WT, *ssh4*Δ, *tul1*Δ, and *ssh4*Δ *tul1*Δ cells. The same volume of cells was loaded, with 0.5 OD_600_ units of cells loaded at 0 h.

Group A contains two substrates: Vph1 and Ypq1. For both proteins, deleting *SSH4* blocked degradation, whereas deleting *TUL1* had no impact on the kinetics (Fig. 5A and S4). In group B, after rapamycin treatment, the degradation of Cot1-GFP was significantly reduced in *tul1*Δ, but nearly unaffected in the *ssh4*Δ strain (Fig. 5B and S4), suggesting that Tul1, but not Ssh4, contributed to the degradation. However, a significant amount of Cot1-GFP was still degraded even in the *ssh4*Δ *tul1*Δ double-deletion strain, indicating the existence of either a new ubiquitin ligase or a new Rsp5 adaptor on the vacuole membrane. Of note, under zinc-depletion conditions, the degradation of Cot1-GFP is mainly dependent on Tul1 and the Dsc complex (Li et al., 2015a), suggesting the action of different recognition mechanisms under these two conditions (i.e., rapamycin vs. Zn^2+^ depletion). Zrt3* and Ypl162c represented the Ssh4-, Tul1-, and unknown E3 ligase-dependent substrates (group C). As shown in Fig. 5C, upon TORC1 inactivation, the degradation of full-length Zrt3*-GFP was partially reduced by either *SSH4* or *TUL1* deletion, and was further decreased, but not completely abolished, in the *ssh4*Δ *tul1*Δ strain, again suggesting the existence of another E3 ligase/Rsp5 adaptor. Similarly, we found that the degradation of Ypl162c also involves an unknown E3 ligase/Rsp5 adaptor, with some contribution from Ssh4 and Tul1 (Fig. 5C).

To directly demonstrate the role of vacuolar E3 ligases in cargo ubiquitination, we performed ubiquitin blots on Vph1 (group A), Cot1 (group B), and Zrt3* (group C). For western detection, the ubiquitin was labeled with a MYC tag. In order to stabilize the ubiquitinated population, we deleted *DOA4* that encodes a major deubiquitinase of the endomembrane system. Because *doa4*Δ *tul1*Δ is lethal (Li et al., 2015a, Tong et al., 2014), we generated a *doa4*Δ *vld1*Δ strain instead to study the role of the vacuolar Dsc complex. Vld1 is a bona fide Dsc component that guides the complex to the vacuole membrane through the AP3 pathway (Yang et al., 2018). Deleting *VLD1* had a similar effect on vacuole membrane degradation as *TUL1* deletion (Fig. 6A, 6C, and 6E).

**Figure 6.**
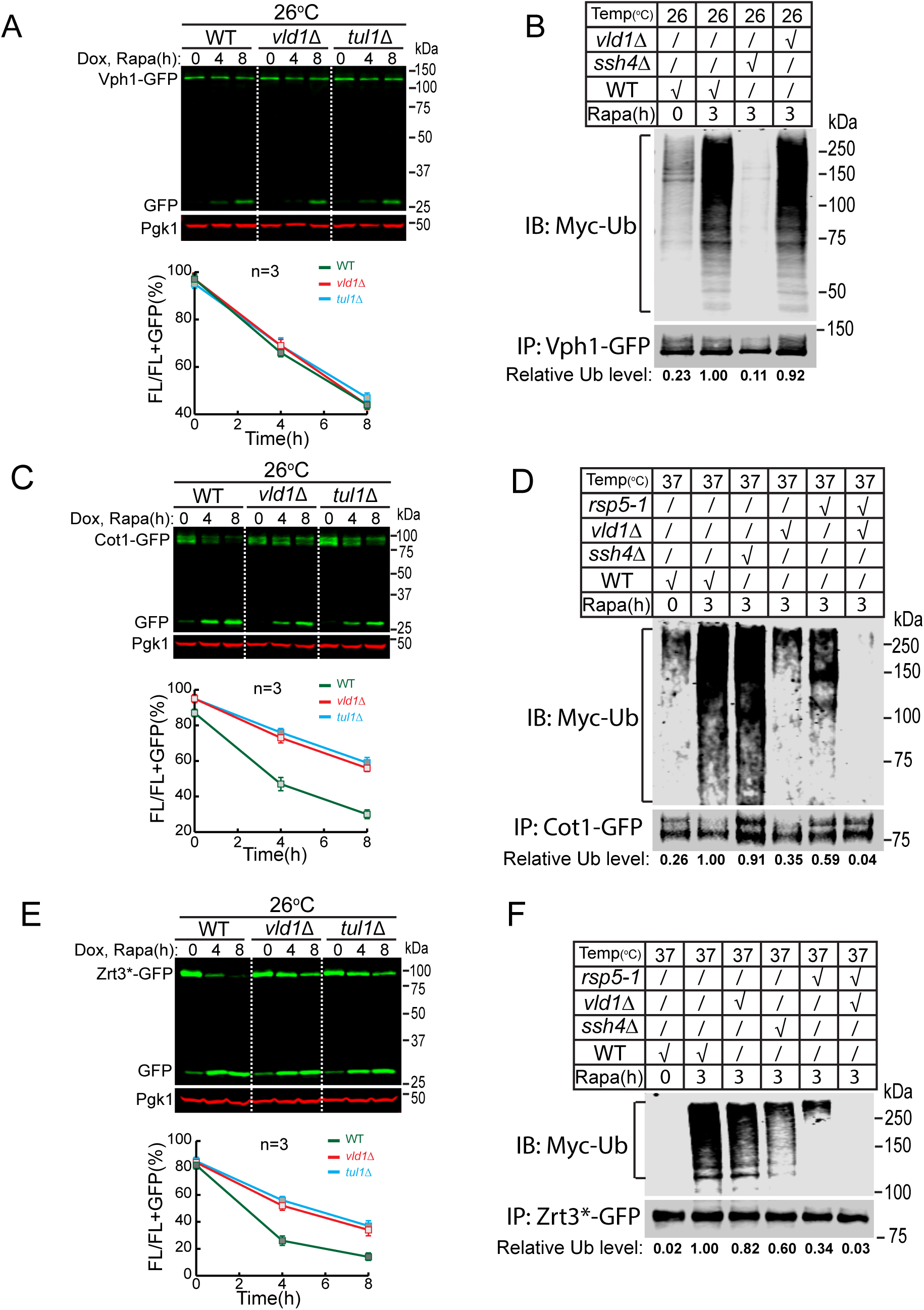
Vacuole membrane E3 ligases poly-ubiquitinate their membrane cargoes upon TORC1 inactivation. **(A)** Western blot (top) and quantification (bottom) showing the degradation of Vph1-GFP in WT, *vld1*Δ, and *tul1*Δ cells. **(B)** A representative western blot (n=3) showing the poly-ubiquitination of Vph1-GFP in WT, *ssh4*Δ, and *vld1*Δ cells at 26°C. The relative ubiquitination level was normalized to the Vph1-GFP level. **(C)** Western blot (top) and quantification (bottom) showing the degradation of Cot1-GFP in WT, *vld1*Δ, and *tul1*Δ cells. **(D)** A representative western blot (n=2) showing the poly-ubiquitination of Cot1-GFP in WT, *ssh4*Δ, *vld1*Δ, *rsp5-1*, and *rsp5-1 vld1*Δ cells at 37°C. **(E)** Western blot (top) and quantification (bottom) showing the degradation of Zrt3*-GFP in WT, *vld1*Δ, and *tul1*Δ cells. **(F)** A representative western blot (n=2) showing the poly-ubiquitination of Zrt3*-GFP in WT, *ssh4*Δ, *vld1*Δ, *rsp5-1*, and *rsp5-1 vld1*Δ cells at 37°C. For (A) (C) and (E), the same volume of cells were loaded, with 0.5 OD_600_ units of cells loaded at 0 h.

As shown in Fig. 6, rapamycin treatment triggered the polyubiquitination of all three proteins. For Vph1, the deletion of *SSH4* eliminated its ubiquitination, whereas deleting *VLD1* had little effect (Fig. 6B, compare the last two lanes). In contrast, *VLD1* deletion led to a reduction (65%) of Cot1 ubiquitination, whereas *SSH4* deletion had little effect (Fig. 6D, lane 3 vs. 4). Interestingly, the temperature-sensitive *rsp5-1* mutation at 37°C resulted in a strong reduction (41%), and further deletion of *VLD1* abolished, the Cot1 ubiquitination (Fig. 6D, lane 5 vs. 6). The different effects between *SSH4* and *RSP5* mutants on Cot1 ubiquitination suggested the existence of a new Rsp5 adaptor, instead of a new E3 ligase. In the case of Zrt3*, deletion of either *SSH4* or *VLD1* caused a reduction (Fig. 6F, 40% and 18%, respectively) of its ubiquitination. However, similar to Cot1, *rsp5-1* mutant had a much stronger reduction (66%) than *ssh4Δ* (18%), and the double mutant of *rsp5-1 vld1*Δ completely abolished the ubiquitination, again suggesting the involvement of a new Rsp5 adaptor.

Our ubiquitin blots suggested the involvement of additional Rsp5 adaptors for Cot1 and Zrt3*. To confirm, we compared their degradation kinetics between the *rsp5-1 tul1*Δ strain and *ssh4*Δ *tul1*Δ strain at a non-permissive temperature. As shown in Fig. 7A-B, the *rsp5-1 tul1*Δ double mutant completely blocked the rapamycin-triggered Cot1-GFP degradation, and no accumulation of free GFP was observed. In contrast, Cot1-GFP was still partially degraded in the *ssh4*Δ *tul1*Δ strain. Therefore, there must be a new Rsp5 adaptor to recognize Cot1.

**Figure 7.**
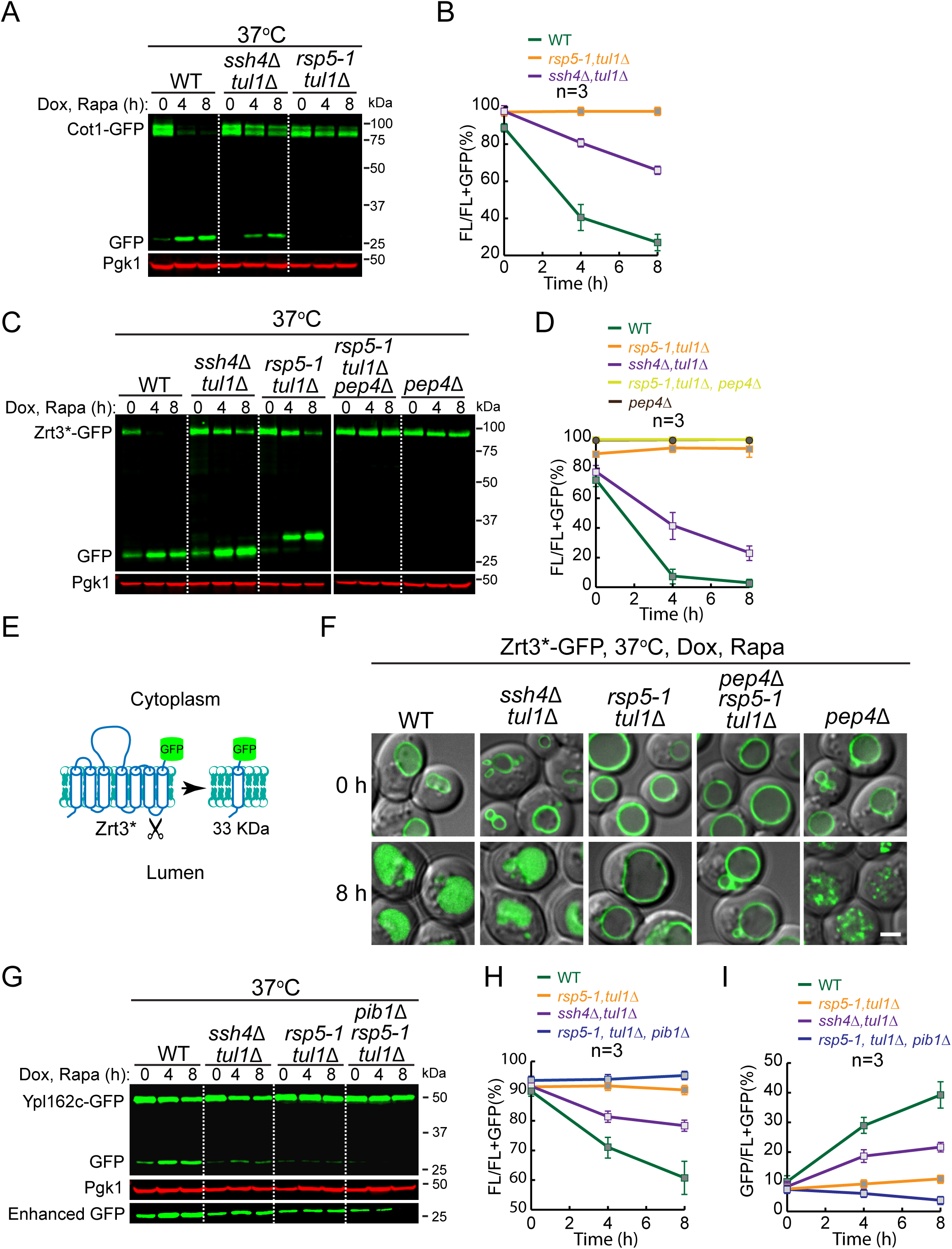
Evidence for the existence of a new Rsp5 adaptor and identification of Pib1 as the third vacuole E3 ligase. **(A)** Western blot showing the degradation of Cot1-GFP in WT, *ssh4*Δ *tul1*Δ, and *rsp5-1 tul1*Δ cells. **(B)** Quantification of the protein levels in (A). **(C)** Western blot showing the degradation of Zrt3*-GFP in WT, *ssh4*Δ *tul1*Δ, *rsp5-1 tul1*Δ, *pep4*Δ *rsp5-1 tul1*Δ, and *pep4*Δ cells. **(D)** Quantification of the protein levels in (C). Please note that the curves of *pep4*Δ *rsp5-1 tul1*Δ and *pep4*Δ samples are almost overlapping. **(E)** A cartoon depicting the cleavage of Zrt3* between TM6 and TM7. **(F)** Subcellular localization of Zrt3*-GFP in the indicated strains before (0 h) and after (8 h) rapamycin treatment. **(G)** Western blot showing the degradation of Ypl162c-GFP in WT, *ssh4*Δ *tul1*Δ, *rsp5-1 tul1*Δ, and *pib1*Δ *rsp5-1 tul1*Δ cells. (**H-I**) Quantification of the FL (H) or free GFP (I) levels in (G). For (A) (C) and (G), the same volume of cells was loaded, with 0.5 OD_600_ units of cells loaded at 0 h. Scale bar: 2 μm.

The degradation of Zrt3*-GFP in the *rsp5-1 tul1*Δ double mutant is very intriguing. As shown in Fig. 7C, the protein levels of the full-length protein were still decreasing in the double mutant at 37°C. However, no increase of free GFP was observed. Instead, an intermediate-sized band (∼33 kDa) accumulated during the rapamycin treatment (Fig. 7C, middle three lanes). These results suggested that the double mutant may have completely blocked the degradation of Zrt3*-GFP. However, Zrt3*-GFP may not be stable at high temperature and was cleaved by a lumenal protease. Based on the size of the cleavage product, the digestion might have happened at the lumenal loop between transmembrane helix 6 and 7 (Fig. 7E). Consistent with this hypothesis, deletion of *PEP4* in the *rsp5-1 tul1*Δ double mutant, or deletion of *PEP4* alone, abolished the accumulation of the 33-kDa band, and no decrease of the full-length protein was observed (Fig. 7C-D). Further supporting evidence was provided by imaging data. As shown in Fig. 7F, no lumenal accumulation of the GFP signal was observed in the *rsp5-1 tul1*Δ double mutant despite the fact that the 33-kDa band accumulated based on the western blot. Importantly, although the *PEP4* single-deletion mutant and the triple mutant displayed a similar phenotype by western blot, they exhibited different phenotypes by fluorescence microscopy. With the triple mutant, Zrt3*-GFP was completely stabilized on the vacuole membrane. In contrast, in the *pep4Δ* mutant, Zrt3*-GFP was detected as small intravacuolar puncta, indicating that it was present on the membrane that had been invaginated, but that the resulting vesicles and their cargoes were not degraded. Together, we concluded that the *rsp5-1 tul1*Δ double mutant completely blocked the ubiquitination of Zrt3*-GFP that was delivered into the vacuole. Furthermore, the difference between the *rsp5-1 tul1*Δ and *ssh4*Δ *tul1*Δ strains also suggested the existence of a new Rsp5 adaptor on the vacuole membrane.

In summary, our analysis indicated that Rsp5 and the Dsc complex are the two major E3 ligases that function downstream of the TORC1 complex to regulate vacuole membrane composition. In addition to Ssh4, there is strong evidence to suggest the existence of another Rsp5 adaptor on the vacuole membrane, which will be characterized and reported elsewhere.

### Identification of a third vacuole E3 ligase, Pib1

Next, we performed the E3 ligase deletion analysis for Ypl162c. Intriguingly, unlike Cot1 and Zrt3*, the degradation of Ypl162c was not completely blocked in the *rsp5-1 tul1*Δ double mutant at 37°C (Fig. 7G-I), indicating the involvement of a third E3 ligase.

To identify the unknown E3 ligase, we generated a triple-mutant strain by further deleting the *PIB1* gene. We focused on Pib1 because this E3 ligase has been localized to the vacuole and endosome membrane but its substrates were so far unknown (Burd and Emr, 1998; Shin et al., 2001). As shown in the last three lanes of Fig. 7G, further deletion of *PIB1* completely abolished the free GFP accumulation. Pib1 is a RING domain-containing E3 ligase with a FYVE domain close to its N terminus (Fig. S5A). As reported, Pib1 is localized to the vacuole membrane and endosomes (Fig. S5B), presumably through its interaction with PtdIns3P (Burd and Emr, 1998; Shin et al., 2001). Furthermore, both ubiquitination and degradation of Ypl162c-GFP were partially reduced in the single *pib1*Δ mutant (Fig. S5C-D), indicating that Pib1 indeed participates in the ubiquitination of Ypl162c. Together, our data suggest that Pib1 plays a role in the regulation of vacuole membrane proteins.

### TORC1 regulates vacuolar E3 ligases

What are the underlying mechanisms for TORC1 to downregulate vacuole membrane proteins? We reasoned that there might be three different levels of regulation: (1) TORC1 may regulate the phosphorylation state of vacuole membrane proteins, (2) TORC1 may regulate the activity of vacuolar ubiquitination machinery, (3) TORC1 may regulate the assembly and disassembly of the ESCRT machinery. Recently, De Virgilio and colleagues reported that TORC1 regulates ESCRT assembly through the phosphorylation of Vps27, a key component of ESCRT-0. Under nutrient-rich conditions, active TORC1 phosphorylates Vps27 and inhibits ESCRT assembly on the vacuole membrane (Hatakeyama et al., 2019). Upon starvation, Vps27 will be dephosphorylated to promote ESCRT assembly. This observation is consistent with the increase of vacuole membrane protein degradation after TORC1 inactivation. However, whether TORC1 regulates the activity of vacuolar E3 ligases is unknown.

To address the relationship between TORC1 and vacuole E3 ligases, we measured the protein levels of the three identified E3 ligase systems after natural starvation. Consistent with the increasing demand for ubiquitinating vacuole membrane proteins, the protein levels of Ssh4 and Pib1 were modestly increased after natural starvation (up to 1.8 fold, Fig. 8A-C). A similar increase was also observed for most components of the Dsc complex, including Ubx3, Tul1, Dsc2, and Dsc3 (Fig. 8D-E). Strikingly, the protein levels of Vld1 were dramatically elevated (6 fold at 12 h, Fig. 8D-E).

**Figure 8.**
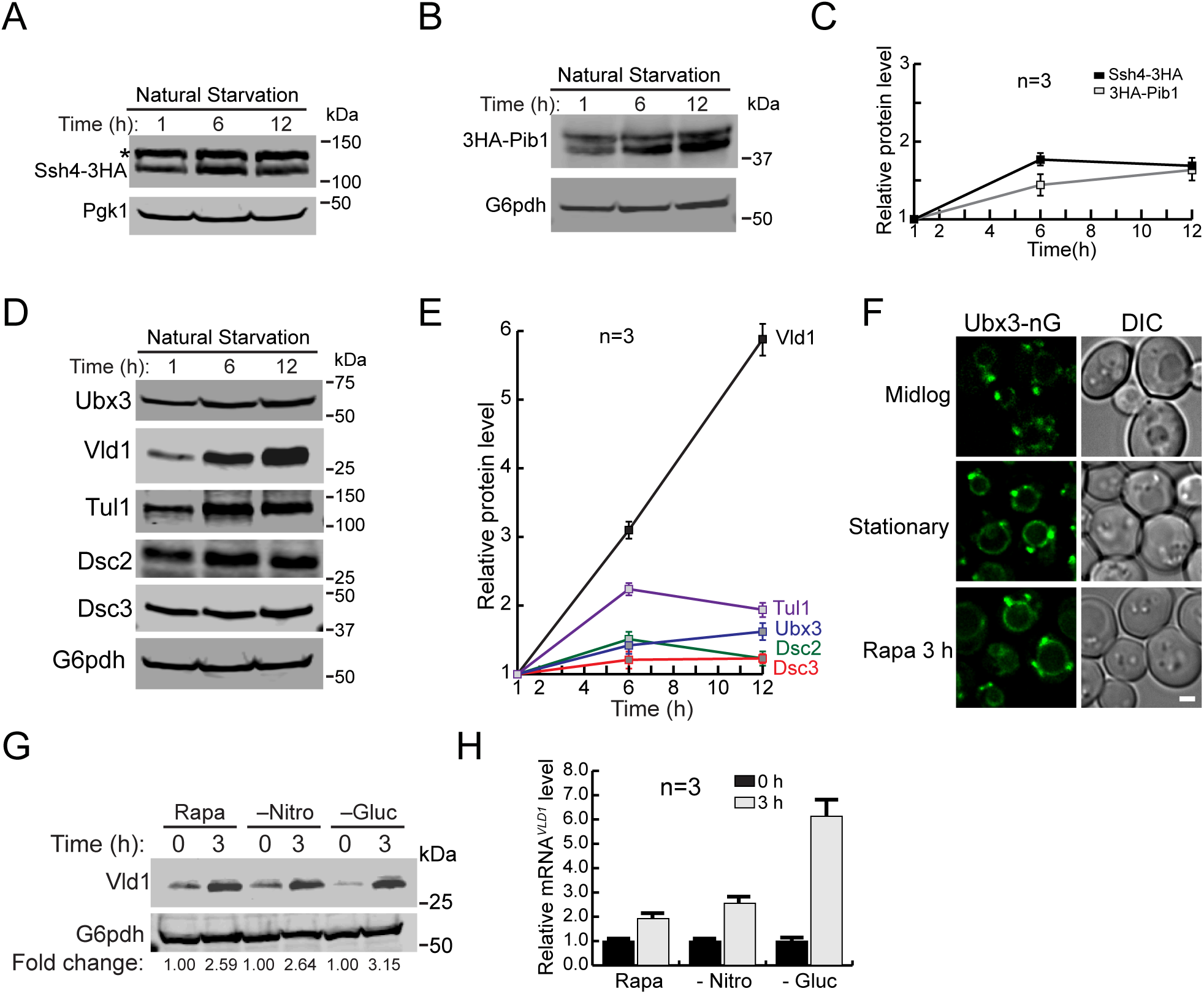
TORC1 regulates the activity of vacuole E3 ligases. **(A)** Western blot showing the protein level changes of Ssh4-mNeonGreen-3HA in stationary phase cells. The asterisk represents a non-specific band. **(B)** Western blot showing the protein level changes of 3HA-Pib1 in stationary phase cells. **(C)** Quantification of the protein levels in (A) and (B). **(D)** Western blots showing the protein level changes of different Dsc components in stationary phase cells. Samples were collected at the indicated time points and 1 OD_600_ units of cells were loaded in each lane. **(E)** Quantification of the protein levels in (D). **(F)** Subcellular localizations of Ubx3-mNeonGreen (Ubx3-nG) in mid-log phase, stationary phase, and rapamycin-treated cells. **(G)** A representative western blot (n=3) showing the level of Vld1-3HA after rapamycin treatment, nitrogen starvation or glucose starvation. Samples were collected at the indicated time points and 1 OD_600_ unit of cells was loaded in each lane. **(H)** qRT-PCR showing the level of *VLD1* mRNA after rapamycin treatment, nitrogen starvation or glucose starvation. Scale bar: 2 μm.

Because Vld1 serves as the vacuole trafficking adaptor of the Dsc complex, its upregulation suggested that TORC1 can regulate the amount of the vacuolar Dsc complex by controlling Vld1 expression. Under nutrient-rich conditions, the Vld1 protein level was low with few vacuole Dsc complexes being assembled (Fig. 8D-F). In contrast, TORC1 inactivation led to the overproduction of Vld1 and increased assembly of the vacuole Dsc complex, as evidenced by the vacuolar localization of Ubx3 (Fig. 8D-F). To verify this model, we tested different conditions that can inactivate TORC1 activity, including rapamycin treatment, nitrogen starvation, and glucose starvation. Strikingly, Vld1 protein upregulation was observed under all conditions (Fig. 8G). As the last test, we measured the *VLD1* mRNA levels using qRT-PCR. As shown in Fig. 8H, the *VLD1* mRNA levels were increased after TORC1 inactivation.

Taken together, we concluded that TORC1 inactivation leads to the upregulation of vacuole E3 ligase systems. Focusing on the Dsc complex, we discovered that TORC1 regulates the amount of the vacuole-localized Dsc (vDsc) complex by controlling the expression of its trafficking adaptor, Vld1.

### TORC1 regulates Vld1 expression through the Rim15-Ume6 signaling cascade

How does TORC1 control the expression of Vld1? The upregulation of *VLD1* mRNA after TORC1 inactivation (Fig. 8H) suggested that it might occur via transcriptional regulation. Using bioinformatics analysis, we identified a URS1 (upstream regulatory site 1) sequence (GGCGGC) ∼500 base pairs upstream of the *VLD1* start codon (Fig. 9A), which is a putative binding site for Ume6. Ume6 is a transcription factor that forms a heterotrimeric complex with Sin3 and Rpd3 (Fig. 9A) (Williams et al., 2002). It has been reported that, under nutrient-rich conditions, Ume6 can repress the expression of *ATG8* by directly binding to the URS1 site of the *ATG8* promoter (Backues et al., 2012; Bartholomew et al., 2012). Importantly, the activity of Ume6 is regulated by TORC1. Upon starvation, Ume6 is phosphorylated through a TORC1-Rim15 cascade to relieve its inhibition of gene transcription. To investigate the importance of Ume6 on *VLD1* transcription, we first examined the *VLD1* mRNA level. In mid-log cells, the *VLD1* mRNA level was ∼ 2-fold higher in the *ume6*Δ strain than the WT strain (Fig. 9B). Furthermore, upregulation of Vld1 protein was detected after deleting either *UME6*, *RPD3*, or *SIN3* (Fig. 9C), indicating their roles in *VLD1* repression. To test the direct binding of Ume6 to the *VLD1* promoter, we applied the chromatin immunoprecipitation (ChIP) analysis using a protein A-tagged Ume6 (Ume6-PA) strain. In this assay, two regions from the *ATG8* promoter served as controls: (a) a region with a confirmed URS1 binding motif (−150) as a positive control, and (b) a region without a binding motif (−700) as a negative control (Fig. 9D). On the *VLD1* promoter, the enrichment of Ume6 was much higher in the URS1 region (−500) than in a region without the URS1 motif (−1000) and was similar to the level of the positive control (Fig. 9D). To further confirm its function with regard to Ume6 binding, we mutated the URS1 motif (GGCGGC to AAAAAA) of the *VLD1* promoter (*VLD1*\*, Fig. 9E) and performed another ChIP analysis. As expected, the URS1 mutation abolished the enrichment of Ume6 on the *VLD1* promoter (Fig. 9E). Taken together, these data suggested that the Ume6 ternary complex suppressed *VLD1* transcription by directly binding to its URS1 motif.

**Figure 9.**
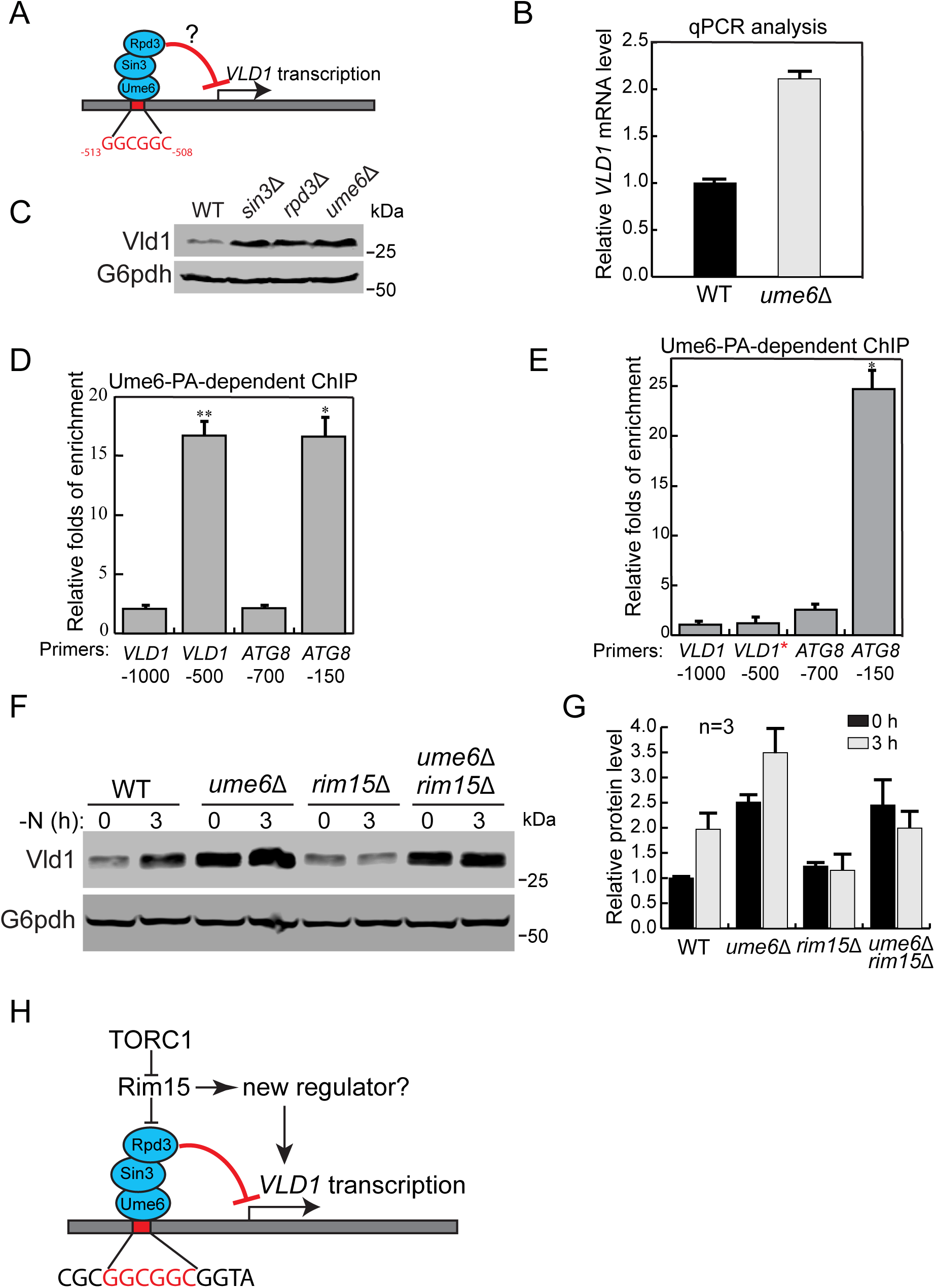
TORC1 regulates Vld1 through the Rim15-Ume6 signaling cascade. **(A)** A cartoon showing the interaction between the Ume6 complex with the putative binding motif in the *VLD1* promoter region. **(B)** qRT-PCR showing the *VLD1* mRNA level in WT and *ume6*Δ cells. **(C)** A representative western blot (n=3) showing the level of Vld1-3HA in WT, *ume6*Δ, *sin3*Δ, and *rpd3*Δ cells. **(D)** ChIP analysis showing the binding of Ume6 to the *VLD1* promoter region. **(E)** ChIP analysis showing the disruption of Ume6 binding to the *VLD1* promoter region after mutation. The enrichment values were normalized to the input DNA, and the error bars show the SEM of 3 independent experiments. The p-value is presented by stars: *<0.05, **<0.01. **(F)** Western blot showing the level of Vld1-3HA in WT, *ume6*Δ, *rim15*Δ, and *ume6*Δ *rim15*Δ cells after nitrogen starvation. Samples were collected at the indicated time points and 1 OD_600_ unit of cells was loaded in each lane. **(G)** Quantification of the protein levels in (F). **(H)** A cartoon model showing that TORC1 regulates Vld1 expression through a Rim15-mediated signaling cascade.

As stated above, Ume6 activity is regulated by TORC1 through the Rim15 kinase (Bartholomew et al., 2012). Based on its phosphorylation state, Rim15 can shuttle between the nucleus and cytosol. Active TORC1 and its downstream effector Sch9 phosphorylate Rim15 and prevent it from entering the nucleus (Wanke et al., 2008; Wanke et al., 2005). In contrast, TORC1 inactivation leads to the dephosphorylation and activation of Rim15 (Pedruzzi et al., 2003). Active Rim15 then enters the nucleus to phosphorylate Ume6 and inhibits its repressor function (Bartholomew et al., 2012). Thus, we asked whether TORC1 is using this signaling cascade to regulate *VLD1* transcription. To this end, we subjected cells to nitrogen starvation to inactivate TORC1 and checked Vld1 protein levels in the WT, *ume6*Δ, *rim15*Δ, and *rim15*Δ *ume6*Δ strains. In WT cells, an increase of Vld1 protein was detected after nitrogen starvation (Fig. 9F-G). In *rim15*Δ cells, the basal level of Vld1 was unchanged; however, there was no upregulation after nitrogen starvation, indicating that Rim15 works downstream of TORC1 as a positive regulator of *VLD1* transcription. Further deletion of *UME6* in the *rim15*Δ strain restored the Vld1 protein level after starvation, suggesting that Rim15 functions upstream of Ume6 (Fig. 9F-G). Interestingly, although the Vld1 protein level was already increased in *ume6*Δ cells before nitrogen starvation, it could be further upregulated after nitrogen starvation (Fig. 9F-G). This result suggested that another regulator besides Ume6 may also function downstream of Rim15 to regulate *VLD1* transcription (Fig. 9H).The identity of this regulator remains to be determined. Nevertheless, our results strongly support the hypothesis that TORC1 uses the Rim15-Ume6 signaling cascade to regulate *VLD1* transcription.

Last, we asked if overexpression of Vld1 alone is sufficient to induce a constitutive degradation of vacuole membrane proteins. Vld1 was overexpressed under the *GPD* promoter and two vacuole membrane cargoes (Cot1 and Zrt3*) were tested. As shown in Fig. 10A-B, before rapamycin treatment, protein levels of both tested substrates were similar between the WT and Vld1 overexpression strains. This indicates Vld1 overexpression alone was not sufficient to trigger a constitutive degradation of vacuole membrane proteins when TORC1 is active. After rapamycin treatment, the degradation kinetics of both Cot1 and Zrt3* were slightly faster upon overexpression. These data support the hypothesis that the TORC1 regulation of vacuole membrane proteins may be controlled at several different levels, and manipulating one condition is not sufficient to induce a dramatic change.

**Figure 10.**
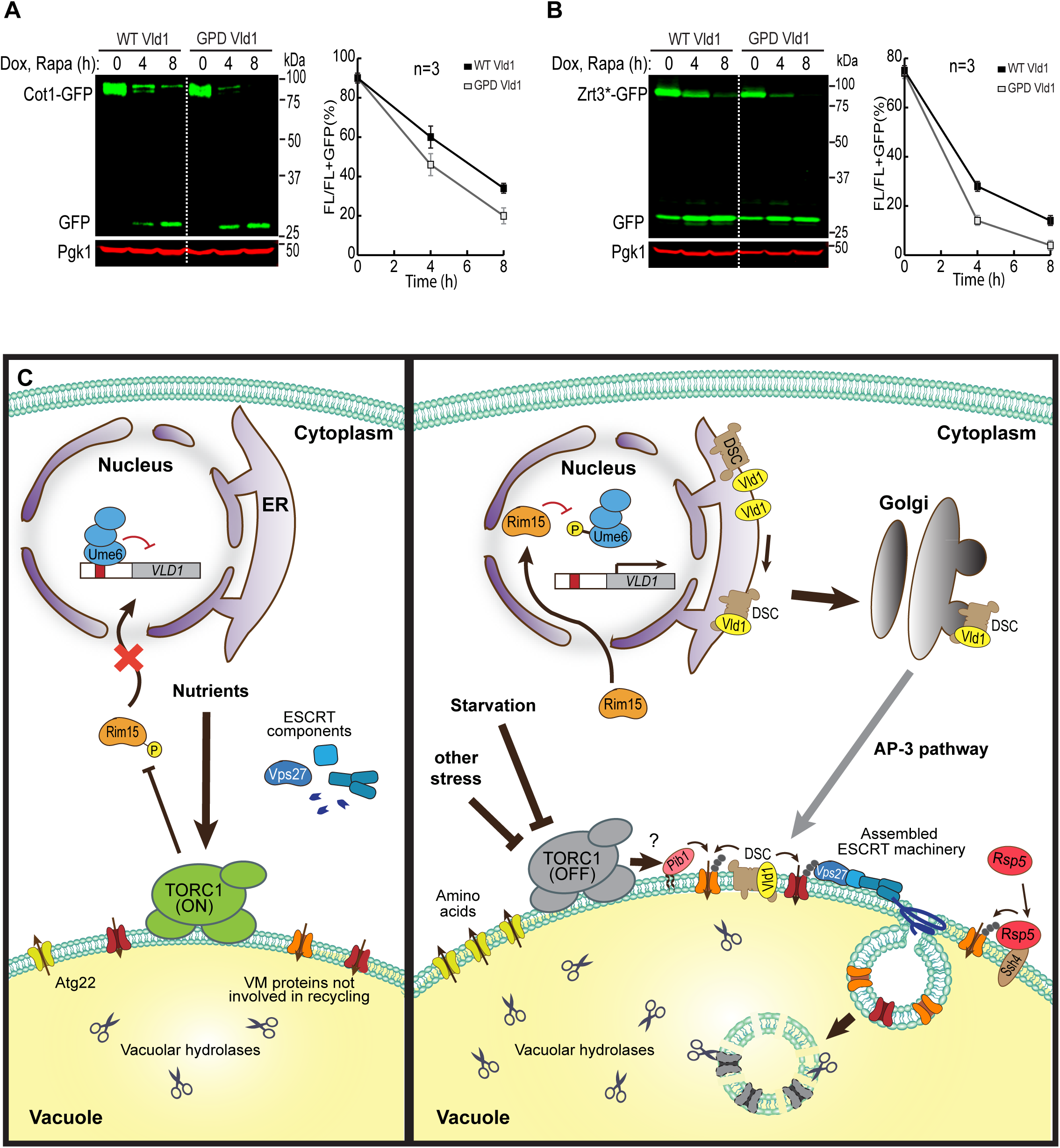
A model summarizing how TORC1 regulates the vacuole membrane composition via the ubiquitin- and ESCRT-dependent microautophagy. **(A-B)** Western blot (left) and quantification (right) showing the degradation of Cot1-GFP (A) or Zrt3*-GFP (B) in WT and Vld1 overexpression strains. The same volume of cells was loaded with 0.5 OD_600_ units of cells loaded at 0 h. **(C)** The model. For details, please see the Discussion.

## Discussion

### TORC1 regulation of the vacuole membrane composition happens at different levels

In this study, we discovered that TORC1 inactivation leads to the downregulation of many vacuole membrane proteins through a ubiquitin- and ESCRT-dependent degradation pathway. This observation is inconsistent with the model that the entire process of vacuole biogenesis will be upregulated upon TORC1 inactivation. We argue that, although lumenal hydrolases and transporters involved in the recycling function are upregulated, many other vacuolar proteins are downregulated to supply additional amino acids for cell survival.

How does TORC1 regulate the vacuole membrane composition? The latter must be controlled at several levels. First, TORC1 can affect the phosphorylation of vacuole membrane proteins. In a large scale proteomic study, Michael Hall and colleagues reported that TORC1 activity can affect the phosphorylation state of vacuole transporters, including Fth1, Ccc1, Avt4, Bpt1, and Fun26 (Soulard et al., 2010). It is conceivable that phosphorylation may prime vacuole membrane proteins for their degradation. Second, as uncovered by our study, under nutrient-rich conditions, active TORC1 inhibits the degradation of vacuole membrane proteins by repressing the activity of ubiquitination machinery. Specifically, active TORC1 inhibits the transcription of *VLD1* through the Rim15-Ume6 cascade. Consequently, very few vDsc complexes can be delivered to the vacuole. When TORC1 is inactive, the inhibition of *VLD1* is relieved, resulting in the upregulation of Vld1 and assembly of more vDsc complexes. By controlling the assembly and trafficking of ubiquitin ligases, TORC1 can regulate the abundance of vacuole membrane proteins in response to environmental cues (Fig. 10C). Third, after ubiquitination, the ESCRT machinery is recruited to the vacuole membrane to sort substrates into the lumen. Interestingly, De Virgilio and colleagues reported that active TORC1 directly phosphorylates Vps27 to inhibit its function on the vacuole membrane. Upon starvation, the dephosphorylation of Vps27 can lead to more efficient assembly of the ESCRT complex on vacuole membrane (Hatakeyama et al., 2019).

In summary, we propose that the vacuole membrane composition is regulated by TORC1 in response to environmental cues. Instead of a simple model that TORC1 inactivation leads to the upregulation of vacuole biogenesis, our study indicated that many membrane proteins are concomitantly degraded to recycle essential amino acids or possibly even “free up” space in the limiting membrane of the vacuole. The regulation may be achieved at three levels, including substrates, E3 ligases, and the ESCRT machinery. We are only at the beginning of understanding this complex relationship.

### Different responses of V-ATPase to the (M)TORC1 inactivation between mammalian and yeast cells

It is intriguing to observe that in yeast, after TORC1 inactivation, Vph1 is downregulated by ∼30-50%. Two other recent publications also made a similar observation (Hatakeyama et al., 2019; Oku et al., 2017). This observation is surprising because, in mammalian cells, it is well established that the v-ATPase components are transcriptionally upregulated by TFEB after MTORC1 inactivation (Sardiello et al., 2009). Considering that the v-ATPase is responsible for vacuole acidification, which is essential for the vacuole’s recycling function (Manolson, et al., 1992), why is it downregulated in yeast?

Three reasons might explain this inconsistency. First, the yeast vacuole pH is maintained at ∼5-5.5(Li and Kane, 2009), which is less acidic than the mammalian lysosome (pH 4.5-5) (Mellman et al., 1986). This difference means the proton concentration inside the vacuole can be up to 10-fold lower than that in the lysosome. Consistent with the pH difference, GFP is quenched by the lower pH and quickly degraded in mammalian lysosomes, whereas in the yeast vacuole, GFP remains fluorescent and resistant to vacuolar proteases. As such, it may require less energy to maintain a proper vacuolar proton concentration. Second, Vph1 is an abundant protein (∼ 20,000 molecules/cell) in yeast (Belle et al., 2006). Accordingly, its partial degradation will not abolish the v-ATPase activity required for maintaining the proton gradient. Instead, this degradation may reduce ATP consumption by the v-ATPase besides supplying extra amino acids for cell survival. Third, consistent with the concept of preserving cellular ATP stores, it is well known that the yeast v-ATPase complex undergoes reversible dissociation between the V_0_ and V_1_ subcomplexes after glucose starvation and other stress conditions (Kane, 1995). In summary, the number of v-ATPase complexes might be more than what is required to maintain a functional vacuole pH when cells are shifted to starvation conditions. Instead, reducing ATP consumption and recycling enough amino acids might be the much more pressing issues for yeast cell survival.

## Experimental Procedures

### Yeast Strains, Plasmids, Media, and Growth Conditions

All yeast strains and plasmids used in this study are listed in Supplementary Table 1. Both Difco^TM^ YPD broth and Difco^TM^ Yeast Nitrogen Base (YNB) w/o Amino Acids and Ammonium Sulfate were purchased from Thermo Fisher Scientific. Yeast Nitrogen Base w/o Amino Acids was purchased from Sigma-Aldrich. All yeast strains were grown at 26°C, unless indicated otherwise, in either YPD or YNB media before further analysis.

### Growth Curve Analysis

Yeast cells were grown in YPD medium to mid-log phase (OD_600_: 0.5 ∼ 0.7) at 28°C, which was arbitrarily defined as time point “1 h” for the growth curve analysis. Then, the growth of the yeast cells was continued at 28°C for up to 40 h. The OD_600_ was measured every 1-2 h and the same number of ODs of cells were collected at the indicated time points for further analysis.

### Rapamycin-triggered Degradation Assay

For substrates that were tagged with GFP or 3xHA at genomic loci, yeast cells were grown in YPD medium to mid-log phase (OD_600_: 0.5 ∼ 0.7), before being incubated with 500 ng/ml rapamycin. After an appropriate amount of time, typically 4-8 h, yeast cells were collected for further analysis. For substrates that were tested under the TET-OFF system, yeast cells were grown in YNB minus uracil medium to mid-log phase (OD_600_: 0.5 ∼ 0.7). The cells were pre-incubated with 2 μg/ml doxycycline for an appropriate amount of time to allow complete ER exit (20 min for most of the substrates, and 1 h for Vph1-GFP, and Fth1-GFP). For the complete ER exit of Fet5-GFP, the plasmid was transformed into a 305-pGpd-Fth1 strain. Yeast cells were then incubated with 500 ng/ml rapamycin for an appropriate amount of time, typically 4-8 h, and collected for further analysis.

### Nitrogen-Starvation Assay

Yeast cells were grown in YPD medium to mid-log phase (OD_600_: 0.5 ∼ 0.7), then pelleted at 3500 rpm for 5 min. After being washed with the nitrogen starvation medium (YNB without amino acids and ammonium sulfate, with 2% glucose) twice, cells were resuspended in the nitrogen starvation medium and incubated at 26°C for an appropriate amount of time (typically 3-4 h). Cells were then collected for further analysis.

### Conventional Transmission Electron Microscopy

Yeast cells were grown in YPD medium to mid-log phase (OD_600_: 0.5 ∼ 0.7), before being incubated with 500 ng/ml rapamycin for 4 h. The samples were further processed in the University of Texas Southwestern Electron Microscopy Core Facility using a published protocol (Hariri et al., 2019; Wright, 2000). Basically, cells were fixed with 2 x prefix solution (4% Glutaraldehyde in 0.2 M PIPES, 0.2 M sorbitol, 2 mM MgCl_2_, 2 mM CaCl_2_), then stained in uranyl acetate and embedded in Spurr Resin. After being polymerized at 60°C overnight, the specimen blocks were sectioned at 70 nm with a diamond knife (Diatome) on a Leica Ultracut UCT 6 ultramicrotome (Leica Microsystems). Sections were poststained with 2% uranyl acetate in water and lead citrate, and were placed on copper grids (Thermo Fisher Scientific). TEM images were acquired on a Tecnai G2 spirit TEM (FEI) equipped with a LaB6 source at 120 kV by using a Gatan Ultrascan charge-coupled device camera.

### Chromatin Immunoprecipitation (ChIP)

ChIP was performed with some modifications from a previously published paper (Aparicio et al., 2005). After the yeast cells were grown to OD_600_ ∼0.8 in YPD medium, formaldehyde was added for DNA-protein cross-linking. Then the DNA was sheared by sonication, and the sheared chromatin was immunoprecipitated. Next, the protein-DNA complex was eluted, and the cross-linking was reversed. Finally, the purified DNA was examined by RT-qPCR analysis. The information for all primers is listed in supplemental Table 2.

### RNA Isolation and qRT-PCR

Total RNA samples were extracted from yeast cells using TRIzol (Life Technologies, 145105) and PureLink^TM^ RNA Mini Kit (Invitrogen, 1938678). For quantitative real-time PCR, approximately 6 μg RNA was applied for 1st-strand cDNA synthesis using PrimeScript^TM^ RT Reagent Kit (TaKaRa, AK6003) with oligo (dT) primers. PCR was then performed using the Power SYBR Green PCR Master Mix (Thermo Fisher, 1708558D) with the primers targeting either *UBC6* (internal control) or specific genes. For each sample, the relative transcript levels were determined by normalizing them to *UBC6* levels. The information for all primers is listed in supplemental Table 2.

### Microscopy and Image Processing

The microscopy and imaging processing was performed with a DeltaVision^TM^ system (GE Healthcare Life Sciences) as described recently in Yang et al., 2018. The filter sets FITC (excitation 475/28, emission 525/48) and TRITC (excitation: 542/27, emission: 594/45), were used for GFP and mCherry, respectively. In brief, yeast cells were washed with milliQ water and imaged immediately at room temperature. Image acquisition and deconvolution were performed with the softWoRx program. The images were further cropped and adjusted by using ImageJ (NIH).

### Immunoprecipitation and Detection of Cargo Ubiquitination

To stabilize ubiquitinated cargoes, the gene encoding the ubiquitin hydrolase Doa4 was deleted in either WT or E3 ligase mutant background. Transient overexpression of MYC-Ub, which was under the control of a copper (*CUP1*)-inducible promoter, was induced by addition of 100 µM Cu_2_SO_4_ for 1 h (2 h for *ssh4*- or *rsp5-1*-related strains) before the cells were treated with rapamycin to trigger cargo ubiquitination. After 3 h of rapamycin treatment in the presence of 100 µM Cu_2_SO_4_, ∼50 OD_600_ units of cells were collected for the immunoprecipitation (IP) experiment.

The IP assay was adapted from Li et al., (2015a), with some modifications. Basically, yeast cells were resuspended in 500 µl IP buffer (50 mM HEPES-KOH, pH 6.8, 150 mM KOAc, 2 mM MgOAc, 1 mM CaCl_2_,15% glycerol) with 0.1% digitonin, supplemented with protease inhibitors and 50 mM n-ethylmalemide. Whole-cell lysates were prepared by bead beating at 4°C for 10 min, followed by addition of 500 µl of 1.9% digitonin in IP buffer. Membranes were then solubilized by nutating lysates at 4°C for 50 min. After removing the pellet by spinning at 13,000g for 10 min, the resulting lysate was incubated with 25 µl GFP-TRAP resin (Chromotek) at 4°C for 1 h. The resin was then washed four times with 0.1% digitonin in IP buffer, and bound proteins were eluted by incubating resin with sample buffer at 65°C for 5 min. The eluates were then analyzed by SDS-PAGE and probed with MYC or GFP antibody.

### Sample Preparation for Western Blotting and Antibodies

Briefly, yeast cells were treated with ice cold 10% trichloroacetic acid (TCA) and incubated on ice for at least 1 h. After washing with 0.1% TCA, the sample pellets were dissolved in 2x boiling buffer (50 mM Tris, pH 7.5, 1 mM EDTA, 1% SDS), disrupted by glass beads using a vortex mixer for 5 min and heated at 65°C for 5 min. After addition of 2x urea sample buffer (150 mM Tris, pH 6.8, 6 M urea, 6% SDS, 40% glycerol, 100 mM DTT, bromophenol blue), samples were mixed by vortex with glass beads for 5 min and incubated at 65°C for another 5 min. The supernatants were collected, subjected to SDS-PAGE and transferred to nitrocellulose membranes for western blotting analysis.

The following antibodies were used in this study: G6PDH (1:10,000; A9521, Sigma), Pgk1 (1:5000; 22C5D8, Invitrogen), mouse anti-GFP (1:500; sc-9996, Santa Cruz Biotechnology, Inc.), rabbit anti-GFP (1:3000; TP401, Torrey Pines Biolabs), anti-HA (1:1000; 16B12, BioLegend), mouse anti-MYC(1:500; 9E10, Santa Cruz Biotechnology, Inc.), rabbit anti-MYC(1:2,000; Sigma), Vph1 (10D7, Invitrogen), Pep4 (1:10,000), Cps1 (1:5,000) (Richter et al., 2007), and Atg8 (1:5,000). Antibodies against Dsc2, Dsc3, Ubx3, and Tul1 were generous gifts from P. Espenshade (Johns Hopkins University, Baltimore, MD).

## Acknowledgments

We thank members of the Li laboratory, including V. Venkatarangan, G. Chu, L. Reist, A. Kappagantu, and A. Hamlin for their technical support. We are also grateful to our colleagues in the Protein Folding and Disease Hub and the MCDB department at the University of Michigan, especially M. Duncan, H. Xu, and Y. Wang for the helpful discussion and critical reading of the manuscript. We thank K. Luby-Phelps and the University of Texas Southwestern Electron Microscopy Core Facility for expert technical assistance. This research is supported by a startup fund and the MCubed 3.0 fund from the University of Michigan and NIH grant GM133873 to M. Li, by NIH grant GM131919 to DJK, and by FWF grants Y444B12, P30263, P29583, and W1101-B18 to DT.

## Author contributions

Conceptualization, X.Y., W.Z., and M.L.; Methodology, X.Y., W.Z., X.W., and M.L.; Investigation, X.Y., W.Z., X.W., P.J.B., D.A.C., S.S, F.M.A., Y.L., and M.L.; Writing – Original Draft, X.Y., W.Z., and X.W.; Writing – Review & Editing, X.Y., W.Z., D.J.K., D.T, and M.L.; Funding Acquisition, Resources, & Supervision, D.J.K., D.T, and M.L.

## Declaration of interests

The authors declare no competing interests.

---

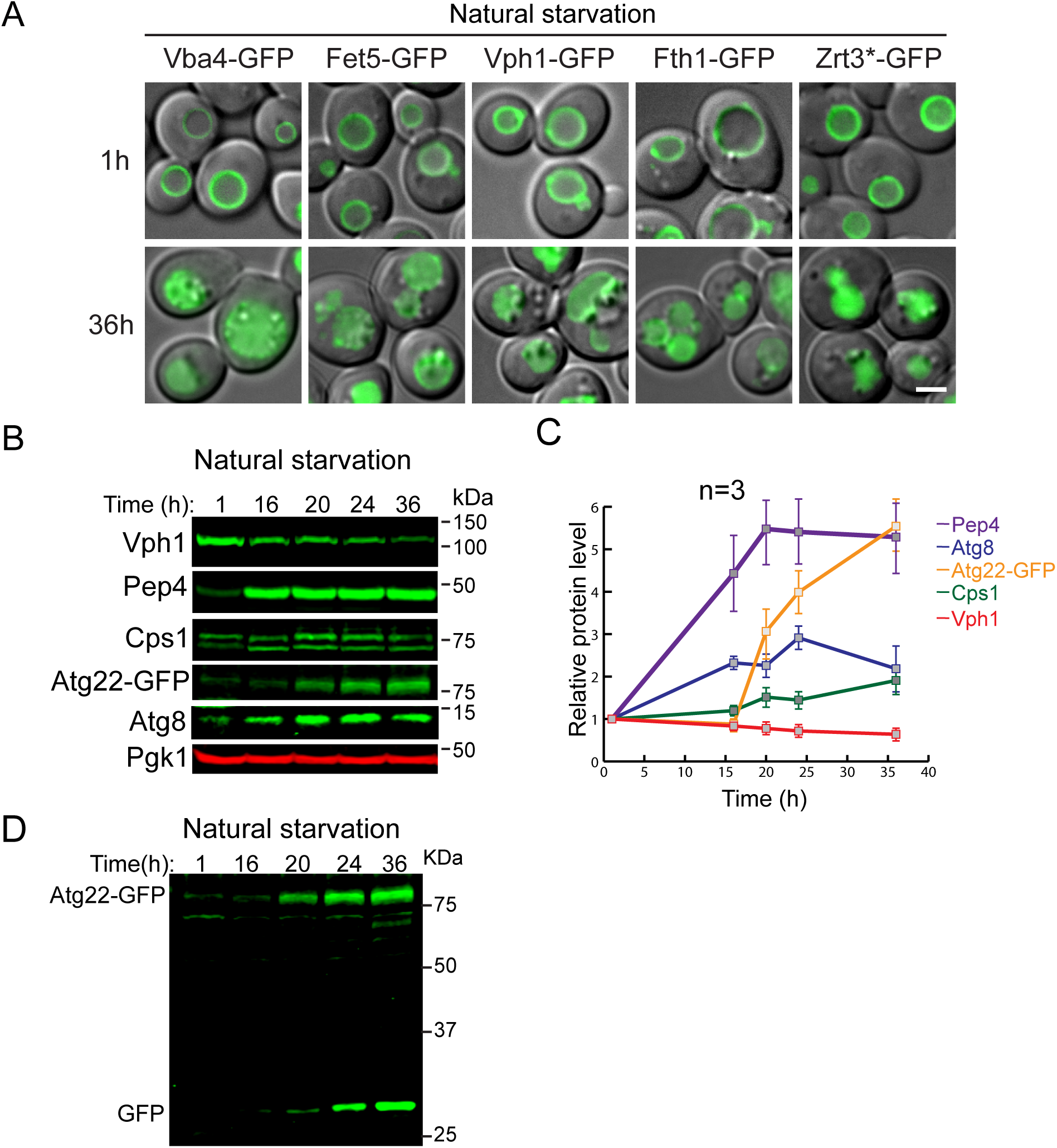

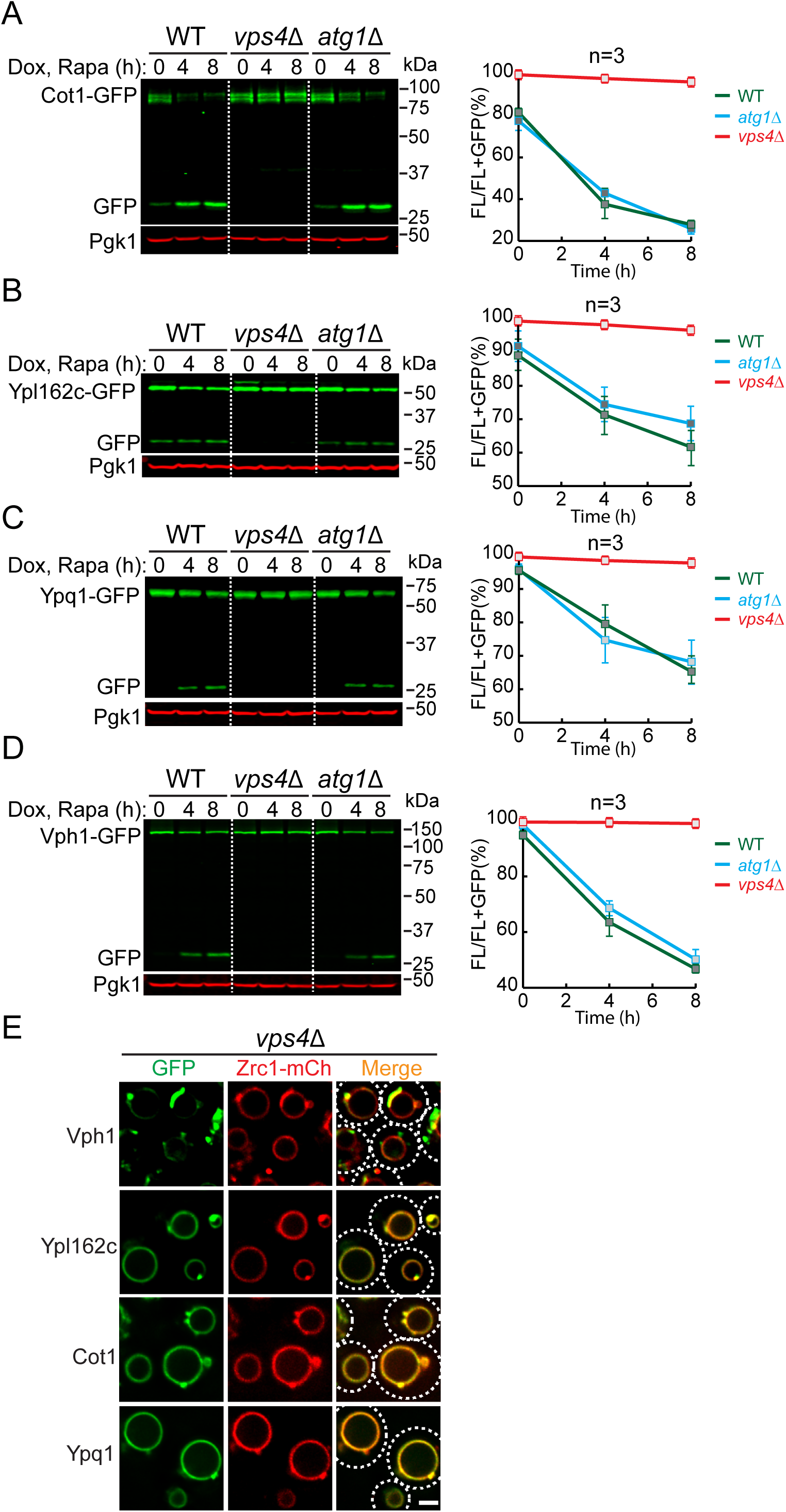

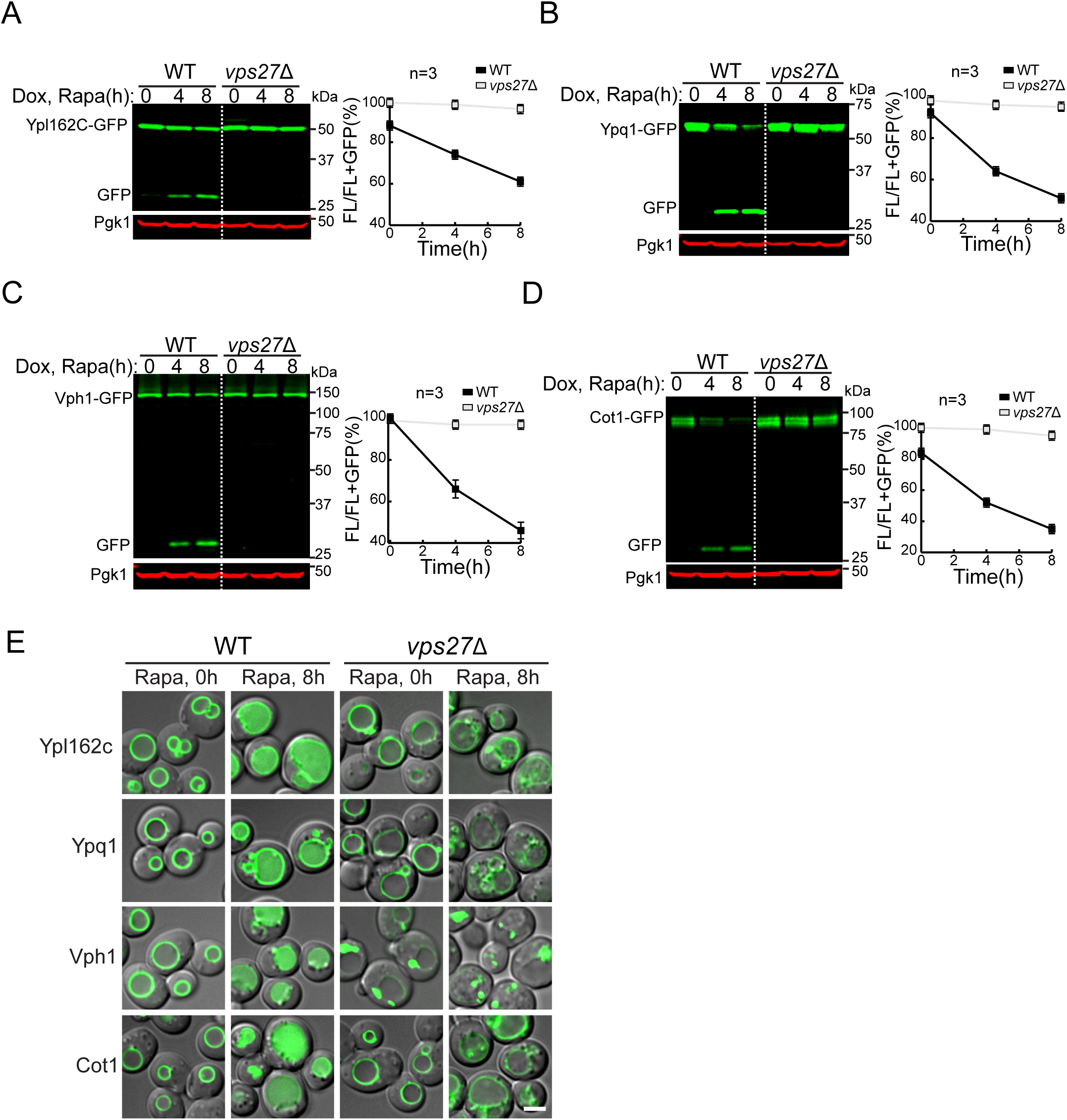

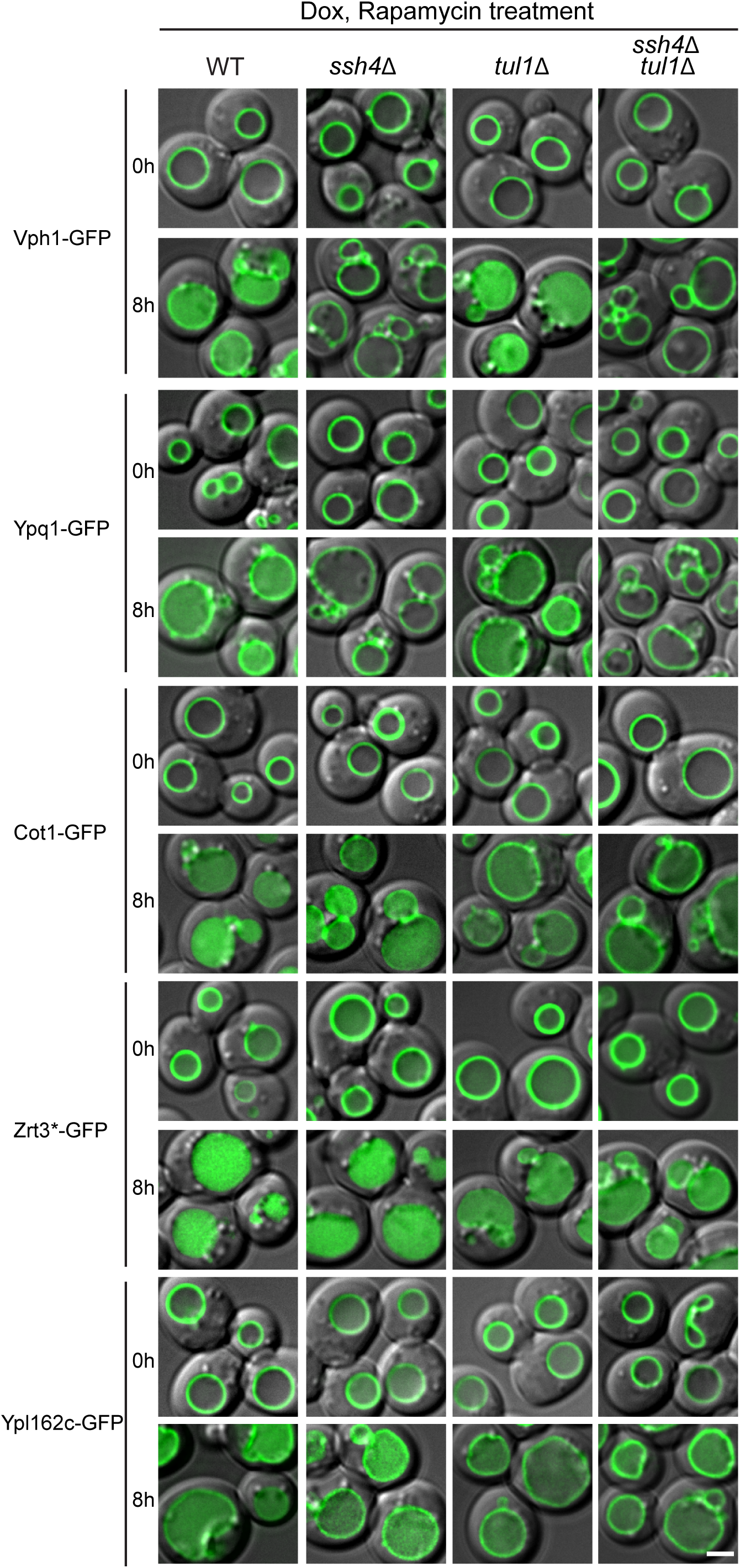

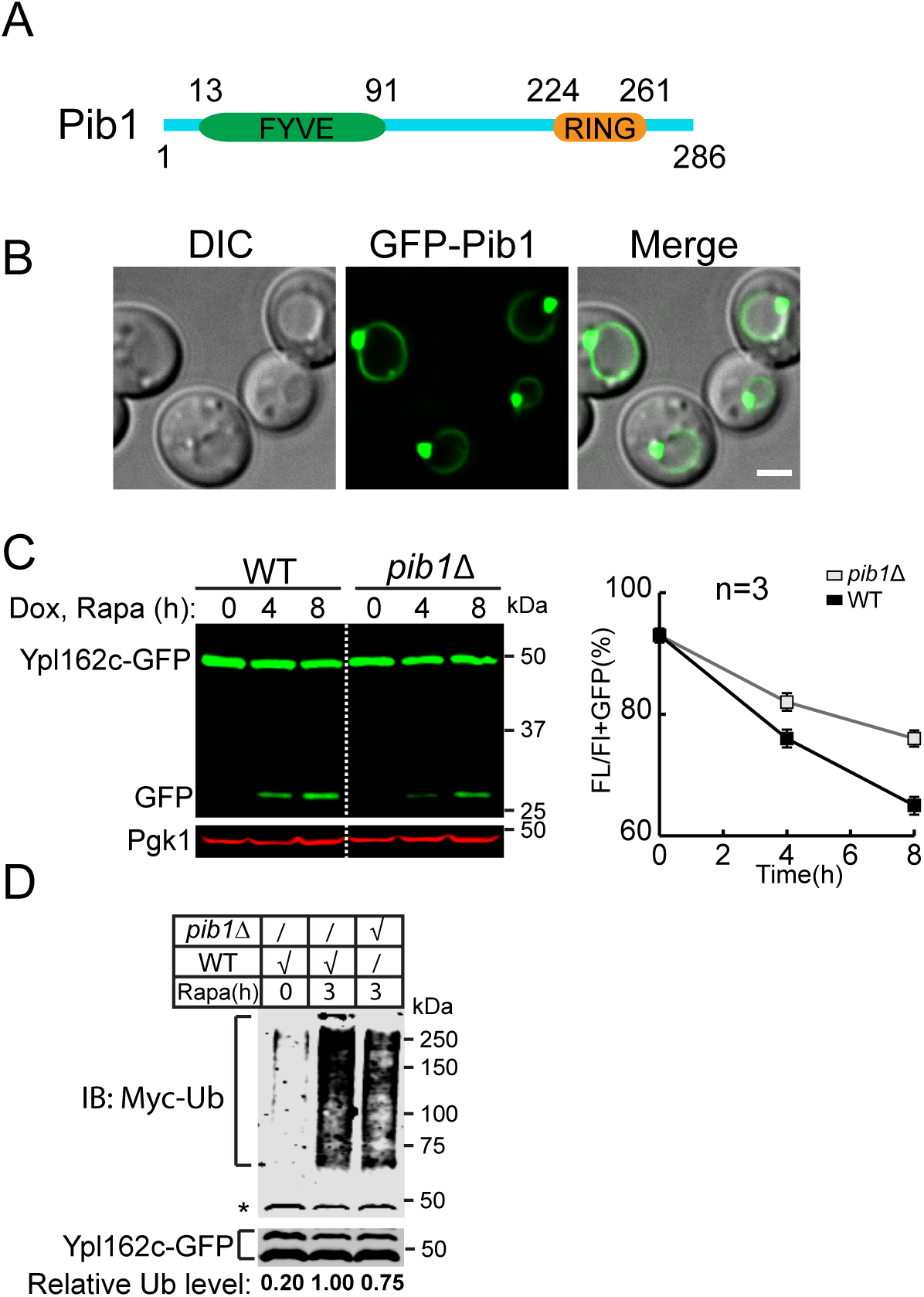

