## Supplemental data for "TORC1 regulates vacuole membrane composition through ubiquitin- and ESCRT-dependent microautophagy"

### Inventory of Supplemental Information

**I. Supplemental Figure Legends**

a. Supplemental Figure 1 Legend

b. Supplemental Figure 2 Legend

c. Supplemental Figure 3 Legend

d. Supplemental Figure 4 Legend

e. Supplemental Figure 5 Legend

**II. Supplemental Table**

a. Supplemental table 1

b. Supplemental table 2

#### I. Supplemental Figure Legends

**Supplemental Figure 1: TORC1 inactivation triggers a global downregulation of vacuole membrane proteins.**

**(A)** Subcellular localization of vacuole membrane proteins in mid-log (1 h) or stationary phase (36 h) cells. **(B)** Western blots showing the changes of Vph1, Pep4, Cps1, Atg8, and Atg22-GFP in stationary phase cells. Samples were collected at the indicated time points and 1 $OD_{600}$ unit of cells was loaded in each lane. **(C)** Quantification of the protein levels in (B). **(D)** Uncropped Atg22-GFP image from (B). Scale bar: 2 μm. Related to figure 1.

**Supplemental Figure 2: ESCRT machinery, but not macroautophagy, is responsible for the degradation of vacuole membrane proteins.**

**(A-D)** Western blots (left) and corresponding quantifications (right) showing the degradation of (A) Cot1-GFP, (B) Ypl162c-GFP, (C) Ypq1-GFP or (D) Vph1-GFP in WT, *vps4*Δ, and *atg1*Δ strain cells. The same volume of cells was loaded, with 0.5 $OD_{600}$ units of cells loaded at 0 h. **(E)** Subcellular localization of vacuole membrane proteins in cells from the *vps4*Δ strain. White dotted circles highlight the yeast cell periphery. Scale bar: 2 μm. Related to figure 3.

**Supplemental Figure 3: ESCRT machinery is responsible for the degradation of vacuole membrane proteins. (A-D)** Western blots (left) and corresponding quantifications (right) showing the degradation of (A) Ypl162c-GFP, (B) Ypq1-GFP, (C) Vph1-GFP or (D) Cot1-GFP in WT and *vps27*Δ strain cells. The same volume of cells was loaded, with 0.5 $OD_{600}$ units of cells loaded at 0 h. **(E)** Subcellular localization of vacuole membrane proteins in WT and *vps27*Δ strain cells before (0 h) or after (8 h) rapamycin treatment. Scale bar: 2 μm. Related to figure 3.

**Supplemental Figure 4: Multiple vacuolar E3 ligases function downstream of the TORC1 kinase.** Subcellular localization of vacuole membrane proteins in WT, *ssh4*Δ, *tul1*Δ, and *ssh4*Δ *tul1*Δ strain cells before (0 h) or after (8 h) rapamycin treatment. Scale bar: 2 μm. Related to figure 5.

**Supplemental Figure 5: Pib1 participates in the ubiquitination. (A)** A schematic model to show the domain organization of Pib1. **(B)** Subcellular localization of GFP-Pib1. **(C)** Western blots (left) and quantification (right) comparing the degradation of Ypl162c-GFP in WT and *pib1*Δ strains. The same volume of cells was loaded, with 0.5 $OD_{600}$ units of cells loaded at 0 h. **(D)** A representative western blot (n=3) showing the poly-ubiquitination of Ypl162c-GFP in WT and *pib1*Δ cells at 28$^{o}$C. The relative ubiquitin level was normalized to the Ypl162c-GFP level. The asterisk represents a background band. Scale bar: 2 μm. Related to figure 7.

#### II. Supplemental Tables

| **Supplemental Table 1: Yeast strains and plasmids used in this study** | | | |
|---|---|---|---|
| ***S. cerevisiae strains*** | | | |
| ***strain*** | ***name*** | ***genotype*** | ***reference/source*** |
| SEY6210 | wild type | Matα*, leu1-3, 112 ura3-52 his3-200, trp1-901 lys2-801 suc2-D9* | (Robinson et al., 1988) |
| SEY6210.1 | wild type | Mata*, leu1-3, 112 ura3-52 his3-200, trp1-901 lys2-801 suc2-D9* | (Robinson et al., 1988) |
| YML235 | Vba4-GFP | 6210, *VBA4-GFP::TRP1* | (Li et al., 2015a) |
| YML106 | Fet5-GFP | 6210.1, *FET5-GFP::HIS3* | (Li et al., 2015a) |
| YML169 | Vph1-GFP | 6210.1, *VPH1-GFP::KAN* | (Li et al., 2015a) |
| YML227 | Zrt3*-GFP | 6210.1, *ZRT3*-GFP::KAN* | (Li et al., 2015a) |
| YML1022 | Ypq1-GFP | 6210, *YPQ1-GFP::TRP1* | (Li et al., 2015a) |
| YML321 | Ypl162c-GFP | 6210.1, *YPL162C-GFP::KAN* | (Li et al., 2015a) |
| YML324 | Cot1-GFP | 6210.1, *COT1-GFP::KAN* | (Li et al., 2015a) |
| YML068 | *vps4*Δ | 6210.1, *vps4*Δ::*TRP1* | (Li et al., 2015a) |
| YML260 | *atg1*Δ | 6210, *atg1*Δ*::KAN* | (Li et al., 2015a) |
| YXY483 | *vps4*Δ, Zrc1-mCh | 6210.1, *vps4*Δ::TRP1, ZRC1-mCherry::HIS3 | This study |
| YML100 | *pep4*Δ | 6210, *pep4Δ*::LEU2 | (Li et al., 2015a) |
| YML208 | *vps4*$^{ts}$ | 6210.1, *vps4Δ*::TRP1, *vps4*$^{ts}$::LEU2 | (Li et al., 2015a) |
| YML354 | *ssh4*Δ | 6210, *ssh4Δ*::TRP1 | (Li et al., 2015a) |
| YML489 | *tul1*Δ | 6210.1, *tul1Δ*::TRP1 | (Li et al., 2015a) |

| | | | |
|---|---|---|---|
| YML505 | *tul1*Δ*, ssh4*Δ | 6210.1, *tul1Δ*::TRP1, *ssh4Δ*::TRP1 | This study |
| YML629 | *rsp5-1, tul1*Δ | 6210, *rsp5-1*, *tul1Δ*::TRP1 | This study |
| YXY464 | *pib1*Δ*, rsp5-1, tul1*Δ | 6210, *pib1Δ*::HYG, *rsp5-1*, *tul1Δ*::TRP1 | This study |
| YXY492 | *pep4*Δ*, rsp5-1, tul1*Δ | 6210, *pib1Δ*::HYG, *rsp5-1*, *tul1Δ*::TRP1 | This study |
| YXY307 | Vld1-3HA,Gld1-3HA | 6210.1, GLD1-3HA::HYG, VLD1-3HA::TRP1 | This study |
| YXY266 | Vld1-3HA | 6210.1, VLD1-3HA::TRP1 | This study |
| YML553 | Ubx3-nG | 6210.1, UBX3-neonGreen::TRP1 | (Yang et al., 2018) |
| YML971 | Ssh4-3HA | 6210, SSH4-NeonGreen-3HA::TRP1 | This study |
| YXY532 | 3HA-Pib1 | 3HA-PIB1::TRP1 | This study |
| YXY331 | Vld1-3HA, *ume6*Δ | 6210.1, *ume6*Δ::HYG,Vld1-3HA::TRP1 | This study |
| YXY333 | Vld1-3HA, *sin3*Δ | 6210.1, *sin3Δ*::HYG,Vld1-3HA::TRP1 | This study |
| YXY334 | Vld1-3HA, *rpd3*Δ | 6210.1, *rpd3Δ*::HYG,Vld1-3HA::TRP1 | This study |
| YXY457 | Ume6-PA | 6210.1, UME6-2PA::Kan, VLD1-3HA::TRP1 | This study |
| YXY458 | Ume6-PA, Vld1* | 6210.1, UME6-2PA::Kan, VLD1$^{mURS}$-3HA::TRP1, | This study |
| YXY332 | Vld1-3HA, *rim15*Δ | 6210.1, *rim15Δ*::HYG,Vld1-3HA::TRP1 | This study |
| YXY440 | Vld1-3HA, *rim15*Δ*, ume6*Δ | 6210.1, *rim15Δ*::HIS3, *ume6Δ*::HYG, Vld1-3HA::TRP1 | This study |
| YXY480 | Fth1 overexpression | 6210.1, 305-pGPD-FTH1 | This study |

| | | | |
|---|---|---|---|
| YML377 | *vps27*Δ | 6210, *vps27*Δ::HIS3 | (Li et al., 2015a) |
| YXY676 | Vph1-GFP, *doa4*Δ | 6210.1, Vph1-GFP::KAN, *doa4*Δ::HIS3 | This study |
| YXY673 | Vph1-GFP, *doa4*Δ, *vld1*Δ | 6210.1, Vph1-GFP::KAN, *vld1*Δ::TRP1, *doa4*Δ::HIS3 | This study |
| YXY675 | Vph1-GFP, *doa4*Δ, *ssh4*Δ | 6210.1, Vph1-GFP::KAN, *ssh4*Δ::TRP1, *doa4*Δ::His3 | This study |
| YXY671 | Cot1-GFP, *doa4*Δ, *ssh4*Δ | 6210.1, Cot1-GFP::KAN, *ssh4*Δ::TRP1, *doa4*Δ::HIS3 | This study |
| YXY672 | Cot1-GFP, *doa4*Δ, *vld1*Δ | 6210.1, Cot1-GFP::KAN, *vld1*Δ::TRP1, *doa4*Δ::HIS3 | This study |
| YXY764 | Cot1-GFP, *doa4*Δ | 6210.1, Cot1-GFP::KAN, ,*doa4*Δ::HIS3 | This study |
| YXY767 | Cot1-GFP, *doa4*Δ, *rsp5-1* | 6210.1, Cot1-GFP::KAN, *rsp5-1*::TRP1, *doa4*Δ::HIS3 | This study |
| YXY768 | Cot1-GFP, *doa4*Δ, *rsp5-1*, *vld1* Δ | 6210.1, Cot1-GFP::KAN, *vld1*Δ::NAT, *rsp5-1*::TRP1, *doa4*Δ::HIS3 | This study |
| YXY755 | Zrt3*-GFP, *doa4*Δ | 6210.1, Zrt3*-GFP::KAN, *doa4*Δ::HIS3 | This study |
| YXY754 | Zrt3*-GFP, *doa4*Δ, *ssh4*Δ | 6210.1, Zrt3*-GFP::KAN, *ssh4*Δ::TRP1, *doa4*Δ::HIS3, | This study |
| YXY753 | Zrt3*-GFP, *doa4*Δ, *vld1*Δ | 6210.1, Zrt3*-GFP::KAN, *vld1*Δ::TRP1, *doa4*Δ::HIS3 | This study |
| YXY785 | Zrt3*-GFP, *doa4*Δ, *rsp5-1* | 6210.1, Zrt3*-GFP::KAN, *rsp5-1*::TRP1, *doa4*Δ::HIS3 | This study |
| YXY786 | Zrt3*-GFP, *doa4*Δ, *rsp5-1*, *vld1* Δ | 6210.1, Zrt3*-GFP::KAN, *vld1*Δ::NAT, *rsp5-1*::TRP1, *doa4*Δ::HIS3 | This study |
| YXY824 | Ypl162c-GFP, *doa4*Δ | 6210.1, Ypl162c-GFP::TRP1, *doa4*Δ::HIS3 | This study |

| | | | |
|---|---|---|---|
| YXY825 | Ypl162c-GFP, *doa4*Δ, *pib1*Δ | 6210.1, Ypl162c-GFP::TRP1, *doa4*Δ::HIS3, *pib1*Δ::HYG | This study |
| YXY489 | *pib1*Δ | 6210.1, *pib1*Δ::HYG | This study |
| MAY143 | Vps4-eGFP | 6210.1, VPS4-3HA-GFP::TRP1 | (Adell et al.,2017) |
| ***S. cerevisiae plasmids*** | | | |
| ***vector*** | ***name*** | ***description*** | ***reference/source*** |
| pCM189 | VBA4-GFP | Tet-off vector, tet-O7+ endogenous promoter C-terminal GFP | This study |
| pCM189 | VPH1-GFP | Tet-off vector, tet-O7+ endogenous promoter C-terminal GFP | This study |
| pCM189 | ZRT3*-GFP | Tet-off vector, tet-O7+ endogenous promoter C-terminal GFP | This study |
| pCM189 | FTH1-GFP | Tet-off vector, tet-O7+ endogenous promoter C-terminal GFP | This study |
| pCM189 | FET5-GFP | Tet-off vector, tet-O7+ endogenous promoter C-terminal GFP | This study |
| pCM189 | ZRC1-GFP | Tet-off vector, tet-O7+ endogenous promoter C-terminal GFP | This study |
| pCM189 | COT1-GFP | Tet-off vector, tet-O7 promoter C-terminal GFP | (Li et al., 2015a) |
| pCM189 | YPQ1-GFP | Tet-off vector, tet-O7+ endogenous promoter C-terminal GFP | This study |
| pCM189 | YPQ2-GFP | Tet-off vector, tet-O7+ endogenous promoter C-terminal GFP | This study |

| | | | |
|---|---|---|---|
| pCM189 | GFP-PIB1 | Tet-off vector, tet-O7 promoter N-terminal GFP | This study |
| pRS425 | pCu-myc-ub | Copper promoter, Myc-tagged Ubiquitin | (Li et al., 2015b) |

| **Supplemental Table 2: Primers used in this study** | | |
|---|---|---|
| ***Primers*** | ***sequence*** | ***purpose*** |
| *VLD1*-500-For | ACTCGATGGCCCTTTCCATGGCAC | ChIP |
| *VLD1*-500-Rev | CAGAGATACTTGAATCTTAGGCACT | ChIP |
| *VLD1*-1000-For | TTTAGCAGCTGCTCTACCGAAGC | ChIP |
| *VLD1*-1000-Rev | GCCTGCTCGACCAAGAACGGGCAT | ChIP |
| *ATG8*-150-For | ATGTAATGCTAACTGTCTCCACC | ChIP |
| *ATG8*-150-Rev | CTCCTCAACCTTTAATGGTTCCC | ChIP |
| *ATG8*-700-For | GTTGGAGGTTGCCGGTATTGA | ChIP |
| *ATG8*-700-Rev | GTCGGTTCTGGTTTCTTGTCC | ChIP |
| *VLD1*-RT-For | AAAGGTCAGTGATAGCGAATTT | qPCR |
| *VLD1*-RT-Rev | AGTACGCTGTTTCTAGAAGTATTAG | qPCR |
| *UBC6*-RT-For | GATACTTGGAATCCTGGCTGGTCTGTCTC | qPCR |
| *UBC6*-RT-Rev | AAAGGGTCTTCTGTTTCATCACCTGTATTTGC | qPCR |
